# Decoding Ethiopia’s unique and structurally divergent *Plasmodium falciparum* populations using genomics

**DOI:** 10.64898/2026.01.19.700308

**Authors:** Ashley A. Osborne, Ashenafi Assefa, Tamiru Shibiru Degaga, Asrat Hailu, Adugna Woyessa, Dagimawie Tadesse, Anjana Rai, Angela Rumaseb, Hidayat Trimarsanto, Dominic Kwiatkowski, Chi Yun Lee, Richard D. Pearson, Ric N. Price, Kamala Thriemer, Sarah Auburn

## Abstract

Previous genome-wide studies suggest that Ethiopian *Plasmodium falciparum* parasites are unique and structurally divergent from sub-Saharan African populations but were limited by small sample sizes. The emergence of molecular markers associated with reduced artemisinin susceptibility in northern Ethiopia, and the drug pressures created by sympatric *P. falciparum* and *P. vivax*, may have fostered a distinct and permissive genetic background for the emergence and spread of multidrug resistant parasites. To characterise parasite population structure and evolutionary dynamics in this setting, we generated whole-genome sequencing data for 163 *P. falciparum* isolates from southern Ethiopia (2017–2021) and integrated these with 854 isolates from across sub-Saharan Africa.

Ethiopian parasites exhibited markedly reduced diversity, highly conserved genomes, and distinct admixture ancestry. Near-fixation of chloroquine and antifolate associated resistance markers, potentially driven by *P. vivax* co-circulation, suggests persistent drug-related selection pressures. We identified a previously undescribed deletion in the *pfmdr1* 5′ UTR that is common in Ethiopia and potentially widespread across African populations. Although clinically relevant *pfk13* variants have emerged in northern Ethiopia, these were not detected in southern Ethiopia. However, strong genetic connectivity between regions indicates the potential for rapid spread of resistance and is therefore important for public health policy and practice.

## Introduction

Despite decades of progress, malaria remains a major public health challenge in sub-Saharan Africa. Driven primarily by the malaria species *Plasmodium falciparum*, the region accounts for 94% of global malaria cases and 95% of deaths^1^. Ethiopia, in particular, faces worrying setbacks on its path to elimination, largely attributable to internal conflicts, the emergence of *Anopheles stephensi* as a competent urban vector, insecticide resistance, financial constraints, and the changing climate^1–3^.

Ethiopia was previously on track to meet its Global Technical Strategy target of a 75% reduction in malaria incidence by 2025, but recent developments have made that goal unattainable and highlight the need for renewed interventions.

The emergence of molecular markers associated with reduced artemisinin susceptibility, most notably the R622I variant within the Kelch13 (*pfk13*) gene reported in northern Ethiopia, threaten treatment efficacy and elimination goals^2,4^. Emerging evidence suggests that R622I frequently arises on genetic backgrounds carrying *pfhrp2* and *pfhrp3* deletions, that are common in Ethiopia and compromise the performance of rapid diagnostic tests (RDTs)^5^. These emerging genetic traits highlight the potential role of parasite population structure in supporting the spread of parasite lineages with adaptive variant combinations. Although artemisinin-based combination therapies (ACTs) remain largely effective, clinical studies have reported concerning trends for delayed parasite clearance following treatment with artemether-lumefantrine (AL), the first-line therapy for most of Africa for more than two decades^3,6,7^.

The horn of Africa is unique compared to most sub-Saharan African regions in that *P. falciparum* and *P. vivax* are sympatric, complicating detection and treatment, as well as introducing nuanced drug pressures onto the parasite populations^5,8^. Although chloroquine is no longer used to treat patients with *P. falciparum* malaria, it remains the first-line treatment for *P. vivax* for a majority of this region^1^. Misdiagnosis and chloroquine’s prolonged half-life result in sub-therapeutic drug concentrations that inadvertently maintain significant chloroquine drug pressure on *P. falciparum* populations. These unique selective pressures on Ethiopian parasites underscore the importance of understanding the complex genetic dynamics of parasites in this region^7,9,10^.

Previous genome-wide studies have identified Ethiopian *P. falciparum* parasites as unique and structurally divergent populations compared with other sub-Saharan African populations^8^. These patterns align with our current understanding of the distinct ancestral origins of the human populations within the Horn of Africa. However, these analyses have been limited by small sample sizes, with only 26 publicly available, high quality, WGS isolates from Ethiopia, precluding temporal resolution or detection of rare adaptive variants. To date, the largest prior investigation used molecular inversion probe (MIP) genotyping, which generated important insights into population connectivity and drug-resistance distribution in northern Ethiopia, but lacked the resolution for haplotype-based analyses of selection^5^.

In this study, we generated whole-genome sequencing (WGS) data from 163 *P. falciparum* isolates collected in Arba Minch, southern Ethiopia, between 2017 and 2021, and integrated these with 53 publicly available isolates from the Horn of Africa and 801 from other sub-Saharan African countries. This combined dataset enabled us to conduct a comprehensive genomic analysis of Ethiopian *P. falciparum* diversity, population structure, and selection. Our results reveal ancestral origins independent from other African lineages, extensive signals of positive selection on drug-resistance loci, and near-fixation of chloroquine-resistance markers. Although the *pfk13* R622I variant was absent in southern Ethiopia, high genomic connectivity between northern and southern populations emphasizes the need for continued genomic surveillance to track the spread of emerging resistance.

## Results and Discussion

### An Africa-wide genomic dataset comprising new Ethiopian P. falciparum genomes

To assess the genomic diversity and variation of southern Ethiopian *P. falciparum* populations, whole genome sequencing (WGS) data was generated from 163 isolates collected in Arba Minch in 2017 (N = 34) and 2021 (N = 129). The WGS data was combined with publicly available Ethiopian data from Oromia collected in 2013 (N = 24) through the Malaria Genomic Epidemiology Consortium (MalariaGEN) Pf7 project^11^. In total, 187 high-quality WGS Ethiopian isolates, comprising 124,133 filtered SNPs, were analysed. For continental comparisons, we combined the Ethiopian data with 830 high-quality Pf7 samples from 13 African countries, producing a final dataset of 1,017 isolates and 861,415 SNPs (Supplementary Table S1). This framework enabled us to evaluate Ethiopian *P. falciparum* population structure, drug resistance, and evolutionary dynamics within an Africa-wide context.

### Comparably low within-host and population diversity in Ethiopia relative to other African regions

The within-host inbreeding coefficient (*F_ws_*) was used to gauge infection complexity. High *F_ws_* values reflect monoclonal infections, while lower values indicate complex infections typical of high transmission settings^12,13^. *F_ws_* estimates were high in Ethiopia (mean *F_ws_* = 0.95; N = 187) compared to Malawi (mean *F_ws_* = 0.68; N = 99) (Supplementary Fig. S1). Across the full dataset, 50.6% of *P. falciparum* isolates were polyclonal (*F_ws_* ≤ 0.95; N = 515), and 49.4% were monoclonal (*F_ws_* ≥ 0.95; N = 502). Ethiopian parasites exhibited the highest genome-wide pairwise identity-by-descent (IBD) across all African populations (mean = 0.23; range = 0.029–1.0; N = 142; *F_ws_*≥ 0.95) (Supplementary Fig. S2–S3), consistent with low recombination, reduced genetic diversity, and limited outcrossing.

There was variation within Ethiopia according to year of sample collection, with the mean IBD ranging from 0.14 in 2013 (range = 0.031 - 1; N = 20), 0.16 in 2017 (range = 0.03 - 0.992; N = 17), and 0.24 in 2021 (range = 0.029 - 1; N = 105).

### High proportion of identity-by-descent maintained over time within Arba Minch P. falciparum genomes

Principal component analysis (PCA) confirmed the limited diversity of Ethiopian isolates relative to other African populations (Supplementary Fig. S4). Unlike identity-by-descent (IBD) metrics, identity-by-state (IBS) metrics do not capture recombination effects^14,15^. To confirm observed trends, IBD measurements were calculated using isolates with high *F_ws_* (≥0.95) to estimate the proportion of genomic regions conserved between the Ethiopian *P. falciparum* genomes across different time points (N = 142), as well as in comparison to other African populations (N = 502).

Closely related individuals tend to contain large and frequent segments of shared regions across the genome, resulting in a high proportion of IBD. In contrast, the genomes of more distantly related individuals, where the genomes undergo regular recombination with genetically distinct individuals, tend to exhibit fewer and smaller shared regions of IBD. These estimates were applied to the parasites genome and calculated by measuring the proportion of pairs identical by descent at each SNP position within the population, where two alleles are defined as identical by state if they have the same nucleotide.

Ethiopian parasites displayed high genomic sharing across time, with most pairs sharing ≥ 25% IBD (Supplementary Fig. S5A). Three isolates from 2013 shared ≥ 50% IBD with those from 2021, indicating the persistence of closely related lineages over nearly a decade. In contrast, the majority of isolates from other African regions exhibited minimal IBD, rarely exceeding 5% (Supplementary Fig. S5B). These trends highlight strong genetic conservation and a restricted parasite gene pool in southern Ethiopia, suggestive of focal transmission and limited recombination.

### Persistence of chloroquine resistance in Ethiopia

The primary biomarker for chloroquine resistance, *pfcrt* K76T, was fixed in Ethiopian isolates (100%; 186/186), in striking contrast to its decline elsewhere in Africa (Supplementary Tables S2-S3; Supplementary File 1)^16^. In other subregions, K76T frequencies ranged from 16.7% in East Africa to 57.6% in Central Africa and 45% in West Africa. Additional *pfcrt* variants, including A220S, Q271E, N326S, and R371I, were nearly fixed in the population. All Ethiopian isolates carried the CVIET haplotype at codons 72, 73, 74, 75, and 76, while the wild-type CVMNK haplotype was absent (Fig. 1A). There was extensive co-occurrence among the variants A220S, N326S, and R371I and the CVIET haplotype. Of the 141 isolates with complete genotype data across all loci (codons 72, 73, 74, 75, 76, 220, 326, and 371), 99.6% (140/141) carried the A220S, N326S, and R371I variants together with the CVIET background. The strong linkage between these positions indicates that they form a dominant *pfcrt* haplotype circulating in southern Ethiopia. Together, with the fixed K76T substitution on the CVIET backbone, this haplotype represents a highly conserved, chloroquine-resistant, lineage. The *pfaat1* S258L variant, characterised to potentiate chloroquine resistance at a reduced fitness cost to the parasite, was also fixed in the Ethiopian population (185/185)^17^.

**Figure 1:**
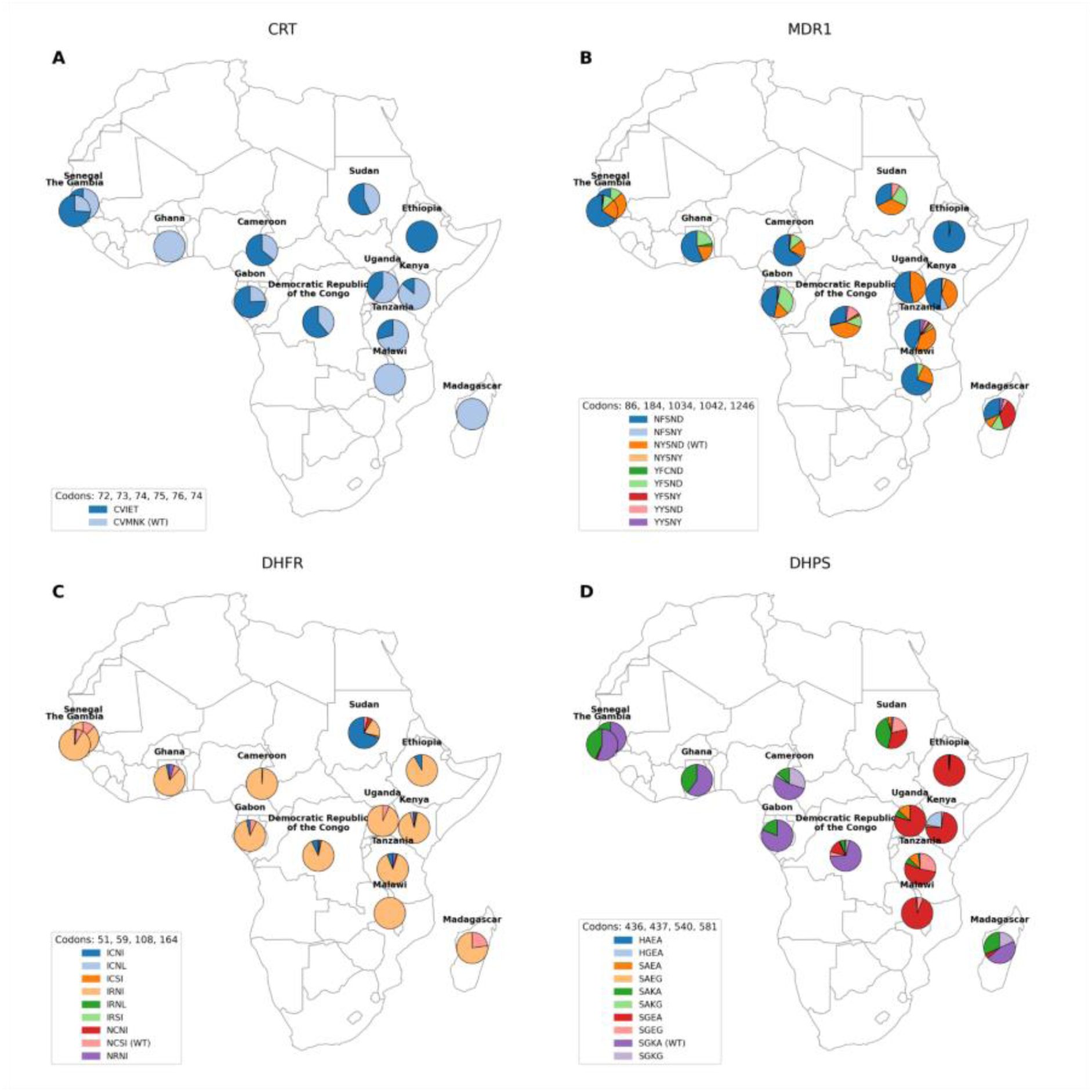
Geographical distribution of resistance haplotypes across sub-Saharan African *P. falciparum* populations on validated drug resistance genes. (A) *pfcrt* codons 72, 72, 74, 75, and 76. (B) *pfmdr1* codons 86, 184, 1034, 1042, and 1246. (C) *pfdhfr* codons 51,59, 108, 164. (D) *pfdhps* codons 436, 437, 540, 581.

Collectively, this suggests strong and sustained selective pressure on *pfcrt* in Ethiopia, likely driven by indirect exposure to chloroquine through treatment of co-endemic *P. vivax*. Misdiagnosis between *P. falciparum* and *P. vivax*, as well as the slow elimination and prolonged subtherapeutic concentrations of chloroquine may have perpetuated the selection of drug-resistant *P. falciparum* parasites^18–21^. Our findings mirror the persistence of chloroquine resistance across northern Ethiopia and underscore the unique evolutionary consequences of *P. vivax – P. falciparum* co-endemicity. The near fixation of *pfcrt* K76T and associated haplotypes demonstrate how ongoing *P. vivax* treatment can indirectly shape *P. falciparum* adaptation, a phenomenon likely to become increasingly relevant as *P. falciparum* declines and non-falciparum malaria species become proportionally more prevalent.

### Antifolate resistance markers in the absence of sulphadoxine-pyrimethamine

Sulphadoxine–pyrimethamine (SP) was discontinued as first line policy in Ethiopia in 2004 and has never been implemented for intermittent preventive treatment in pregnancy (IPTp-SP) or seasonal malaria chemoprevention (SMC). Despite a prolonged period of low drug pressure, antifolate resistance markers persist at exceptionally high frequencies in our dataset. The *pfdhfr* triple mutant IRNI haplotype (codons 51, 59, 108, 164) was predominant (90.9%; 169/186), and the double mutant ICNI haplotype accounted for the remaining isolates (9.1%; 17/186) (Fig. 1C). The wild-type NCSI haplotype was absent. Similarly, the *pfdhps* SGEA double mutant haplotype (codons 436, 437, 540, 581) was observed in 98.4% (183/186) of isolates, followed by the WT SAKA haplotype (1.1%; 2/186) and SGEG triple mutant (0.5%; 1/186). The K540E mutation, characterized as a strong modulator of resistance and decreased efficacy of IPTp-SP, was present in nearly all Ethiopian samples (98.9%; 183/186) but only present in 1% (2/197) of WAF isolates (Supp. Table S3)^22^. These frequencies are comparable to, or exceed, those in regions where SP is still in active use (Fig. 1C, D; Supp. Table S3). The persistence of antifolate-resistant haplotypes may reflect low fitness costs combined with secondary drug pressure from co-trimoxazole, an antibiotic with a similar target pathway, used widely for bacterial prophylaxis and treatment in Ethiopia^23,24^.

### Absence of pfk13 artemisinin-resistance variants despite high IBD connectivity

Artemisinin partial resistance has been reported in other regions of Ethiopia, no *pfk13* variants associated with partial artemisinin resistance were detected in Arba Minch isolates, including the emergent R622I or P441L, A578S, and A675V variants^4^. Only three variants, D89E, K189T, and Q234, were identified at very low frequencies (Supp. Table S3). Previous studies suggest that *pfmdr1* and *pfcrt* mutations may be linked to a genetic profile with a high affinity for acquiring further drug resistance, including resistance to artemisinin, with the N326S variant on *pfcrt* implicated in resistance to artemisinin in Southeast Asia^7,25,26^. The *pfmdr1* NFSND haplotype (codons 86, 184, 1034, 1042, and 1246) dominated (98.4%; 183/186), while the wild-type NYSND haplotype was absent in Ethiopian populations despite being present in all other African populations (Fig. 1B). The NFSND-haplotype has been implicated in reduced efficacy of both lumefantrine and chloroquine^4,7,27–29^. Recent evidence suggests that variants on the *pfPX1* locus (L1222P, M1701I and D1705N) may be associated with decreased susceptibility of *P. falciparum* to lumefantrine and dihydroartemisinin. None of these variants were detected in our Ethiopian dataset and only L1222P was detected at low frequencies in our wider African dataset (Supplementary Tables S3)^30^.

Comparison of Arba Minch WGS data with molecular inversion probe (MIP) data from 525 isolates (SNPs = 1,034) collected across Amhara, Gambella, Oromia, and Tigray revealed extensive connectivity, with numerous southern–northern isolate pairs sharing ≥ 50% IBD and some ≥ 95% IBD (Fig. 2)^5^. Despite this connectivity, the absence of *pfk13* resistance alleles in southern Ethiopia suggests that resistance has not yet spread from northern populations, providing an opportunity for containment. Ongoing molecular surveillance in southern regions will be vital for tracking the extent to which these variants have spread.

**Figure 2:**
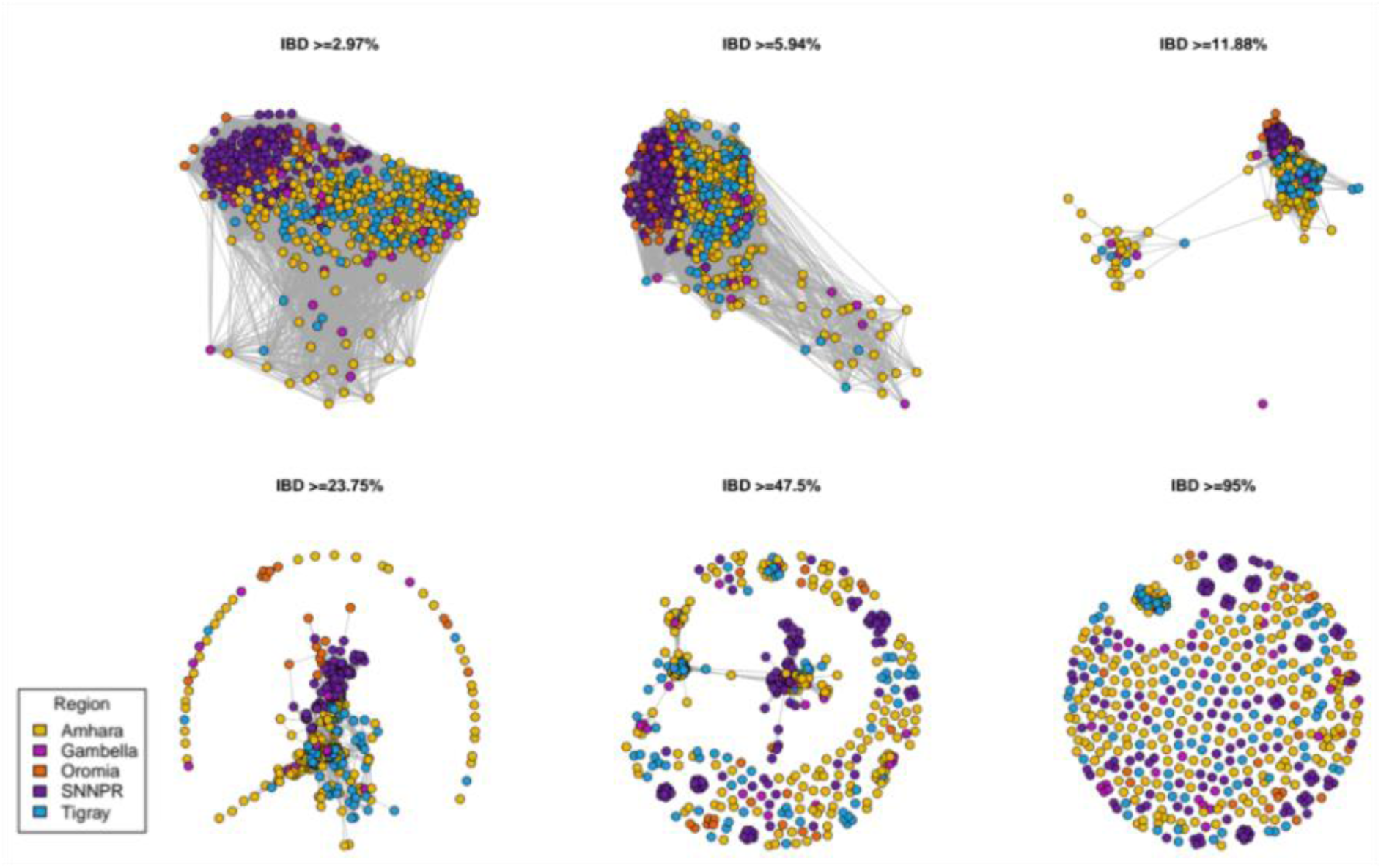
Identity-by-Descent connectivity plot of Ethiopian isolates from Arba Minch (SNNPR) alongside Molecular Inversion Probe data from other regions within Ethiopia (N = 525; SNPs = 1,034).

### Structural variants detected on key diagnostic and drug resistance loci

Structural variant analysis of *P. falciparum* isolates from southern Ethiopia identified total loss of coverage affecting the *pfhrp2* and *pfhrp3* loci in a large proportion of samples, as well as a previously undescribed microdeletion within the 5′ untranslated region (UTR) of *pfmdr1* (Supplementary Tables S4-S5; Supplementary Figure S6).

Across 156 Ethiopian isolates, 5.12% (8/156) exhibited evidence of a complete loss of coverage across *pfhrp2* starting at ∼1.3745 Mbp (Supplementary Table S4). All isolates carrying this deletion were collected in 2021 (7.41%; N = 8/108). Out of 169 isolates, 94 exhibited a total loss of coverage across *pfhrp3* from either a breakpoint around 2.8075 Mbs (2013 [42.9%; N = 9/21] and 2021 [56.4%; N = 57/101]) or 2.8065 Mbp (2017 [59.6%; N = 28/47]). Of the 8 samples that had a loss of *pfhrp2*, 7 retained coverage of *pfhrp3* and the remaining isolate had poor sequencing quality preceding the genomic region of interest. One, or both, of these losses in coverage over these genes may impact the diagnostic sequence relevant to HRP2/3-based rapid diagnostic test (RDT) detection and impact sensitivity^1,5,31^. These findings align with previous descriptions demonstrating that *pfhrp3* deletions are now widespread across the Horn of Africa and concerning reports that these deletions may precede the subsequent loss of *pfhrp2*. Recent findings in Ethiopia also suggest that deletions in *pfhrp2* and *pfhrp3* may be linked with the acquisition of the *pfk13* variant R622I, potentially complicating not only diagnosis but also effective treatment of *P. falciparum* in these regions^5^.

Analysis of *pfmdr1* revealed a 22–36 bp indel, or microdeletion, within the 5′ UTR in 40% (26/65) Ethiopian isolates that passed quality control thresholds, including a median regional depth ≥25 and ≥90% regional coverage at a depth of ≥20. (Supplementary Table S5 and Figure S6). The prevalence of this polymorphism in Ethiopia stayed at 40% when considering only samples with an *F_ws_* ≥ 0.95. This rose to 43.1% (25/58), or 42.9% (*F_ws_* ≥ 0.95; N = 22/55), when considering samples only from 2021. The same indel was present in 32.2% (19/59) and 34.5% (10/29; *F_ws_* ≥ 0.95) of East African isolates (Kenya, Tanzania, and Uganda) and in 32.6% (30/92) and 22.9% (8/35; *F_ws_* ≥ 0.95) of Central African isolates (Cameroon and Gabon) under identical filtering criteria.

A large proportion of isolates had insufficient regional coverage of the *pfmdr1* region to be included in this analysis without the risk of false positive detection, resulting in small sample sizes. Long-read deep sequencing may provide more accurate prevalence estimates and increase the detection power for this variant. Given the indels position within the 5′ UTR of *pfmdr1*, this variant may influence transcriptional regulation and has the potential to modulate a mechanism of drug resistance, as hypothesized for *P. vivax* populations in Cambodia^32^. This microdeletion represents a previously undescribed indel in African *P. falciparum* populations and may highlight ongoing diversification at both drug-response and diagnostic loci. Another important consideration when estimating the prevalence of this variant is that common amplicon-based sequencing panels do not tent to encompass the 5’ UTR of *pfmdr1*, resulting in a lack of representative sequencing data for comparison and screening purposes.

### Temporal and cross-population signals of selection

We hypothesised that prolonged artemisinin, partner drug, and residual chloroquine exposure would result in temporal shifts in selection signals. To capture genomic regions undergoing selection within Ethiopia, haplotype structure was assessed using the integrated haplotype score (*iHS*), which determines levels of homozygosity across the genome. SNPs in regions of the genome exhibiting an *iHS* value of ≥2.5 or ≤-2.5 were classified as significant. This analysis identified a number of regions within Ethiopian populations (*F_ws_* ≥0.95; N = 502) undergoing selection across years, particularly within genes linked to host immune evasion (pfemp1, cytoadherence proteins, and conserved exported proteins) (Supplementary Fig. S7; Supplementary File 2)^12,33^.

We observed repeated *iHS* signals in invasion and cytoadherence genes, with CLAG3.1 and CLAG3.2 significant in 2013, 2017, and 2021 (CLAG3.1: 26 cumulative hits; CLAG3.2: 16). We also replicated a *pfUT* signal in 2013 (max *iHS* = 2.911) and 2017 (iHS = 2.546), with *pfUT* hypothesized to be involved in resistance to quinines and quinidine, although no direct link has been confirmed from *ex vivo* drug susceptibility experiments^34^. Antifolate selection peaked in 2013 at *pfdhps* (PF3D7_0810800 position 548801; *iHS* = 2.91) with attenuation in later years, aligning with haplotype fixation and diminished within-population contrast. Our results highlight a recurring signature of selection in genes mediating invasion, transport, and immune evasion, alongside loci under established drug pressure.

In addition to *iHS* scores, regions exhibiting excessive proportions of IBD, compared to the surrounding genome, were assessed as another proxy for selection. Regions with an IBD fraction above the 90th quantile of values across the genome were classified as significant (Figure 3; Supplementary Figure S8; Supplementary File 2). IBD-based selection scans revealed recurrent hotspots across validated drug resistance associated genes, including *pfmdr1, pfcrt, pfdhfr,* and *pfdhps*. Other drug resistance candidate genes were also identified to exhibit excessive proportions of IBD, including *pfaat1*, *pfap2mu*, and *pfgch1,* with *pfap2mu* hypothesized to be involved in artemisinin resistance (chr12: 700-750 kb; IBD = 0.207 [2017]) and *pfgch1* potentially implicated in antifolate resistance (chr12: 950–1000 kb; IBD = 0.161 [2013] / 0.200 [2017] / 0.308 [2021]). In both 2013 and 2017 isolates, high IBD regions surrounding *pfdhps* were observed (chr8: 500–550 kb; IBD = 0.322 [2013] / 0.369 [2017] and chr8: 550-600 kb; IBD = 0.277 [2013] / 0.305 [2017]), while *pfdhfr* was only observed in 2013 and 2021 isolates (chr4: 700–750 kb; IBD = 0.163 [2013] / 0.339 [2021]).

**Figure 3:**
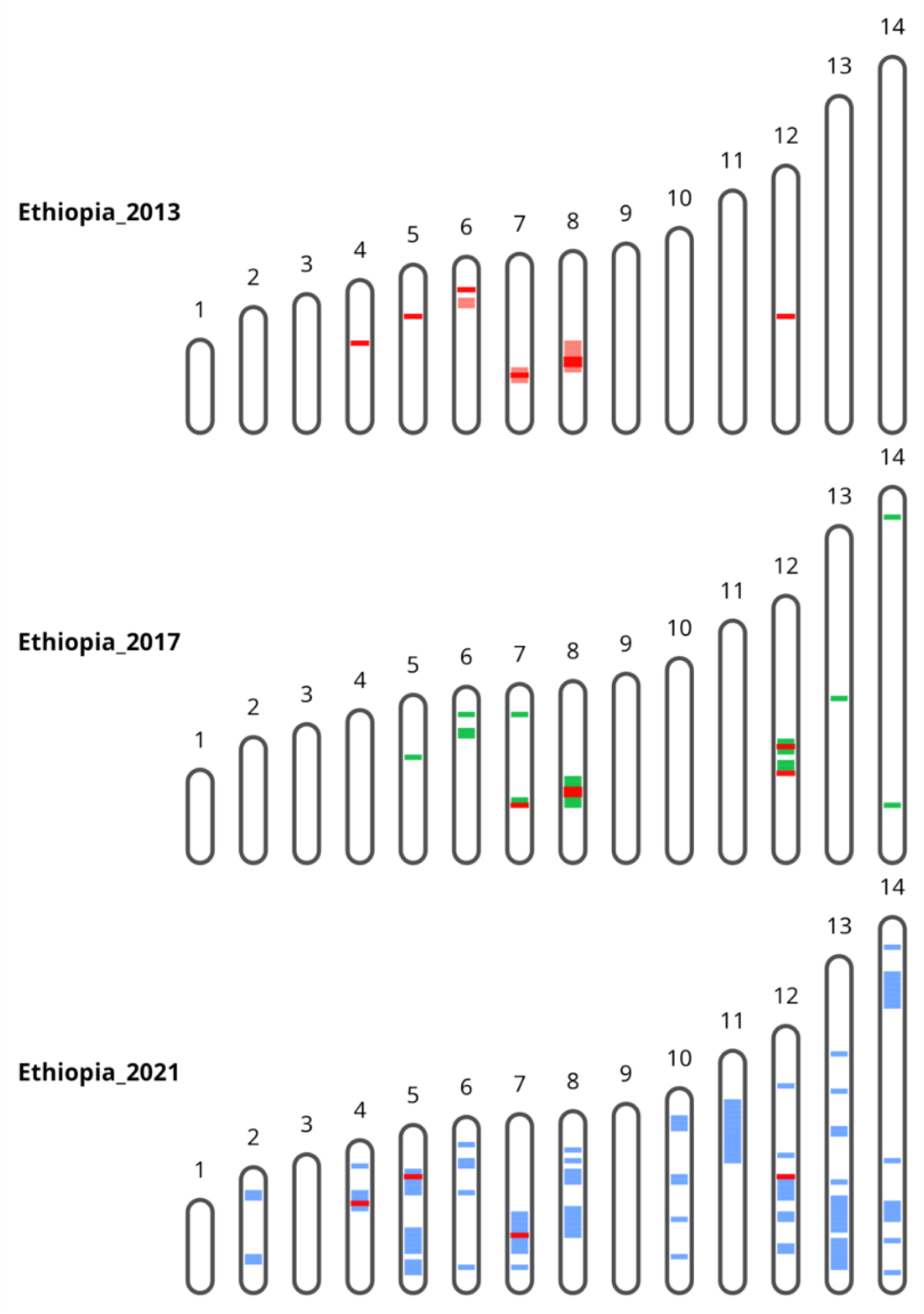
Chromosome map of regions exhibiting high fractions of Identity-by-Descent (90th quantile) over time in Ethiopia. Regions containing genes associated with drug resistance (Supplementary Table S2) are highlighted in red while region colour corresponds to year of collection* (window size 50kb). *2013, N = 20; 2017, N = 17; 2021 = 105

Despite ongoing chloroquine pressure within the region, and the near fixation of molecular markers associated with resistance, high fractions of IBD on *pfcrt* (chr7: 400–450 kb; IBD = 0.431 [2013] / 0.468 [2017] / 0.324 [2021]) were observed in all Ethiopian isolates, consistent with historic rather than ongoing chloroquine selection. High fractions of IBD proportions were also identified on *pfaat1* in isolated from 2013 (chr6: 1200–1250 kb; IBD = 0.199 [2013]). In the 2013 and 2021 parasite populations, regions encompassing the *pfmdr1* gene, a known modulator of resistance to chloroquine and lumefantrine, exhibited excessive IBD (chr5: 950–1000 kb; IBD = 0.171 [2013] / 0.328 [2021])^3,28,35^. Recent evidence suggests specific variations in *pfmdr1* may support the acquisition of additional variants conferring resistance to artemisinin, including *pfk13* variants which are rapidly expanding in Ethiopia^36^. Beyond classical resistance targets, single-window peaks included AP2-G, SET10, HP1, and PTEX88, regulators of commitment, chromatin remodelling, and protein export, suggesting broader adaptive responses to host and environmental pressures alongside drug-driven evolution.

Cross-population extended haplotype homozygosity (*XP-EHH*) was calculated for isolates from across the African continent (*F_ws_* ≥0.95; N = 502) (Supplementary Fig. S9; Supplementary File 2). *XP-EHH* values ≥ 2.5 and ≤ -2.5 were considered significant. This analysis identified strong differentiation at *pfmrp1* (*XP-EHH* = −2.51 to −2.54) in Ethiopia relative to populations in Ghana and Senegal, possibly indicating divergent selection on multidrug-resistance pathways^37^. Immune-associated loci featured among observed extreme values, including MSP7 (Ethiopia vs Uganda; *XP-EHH* = 3.88 to 4.50), reinforcing parallel selection on invasion and cytoadherence, as well as multidrug-resistance mechanisms.

### Genetic differentiation and distinct ancestry of Ethiopian populations

To identify alleles under long-term or region-specific selection, fixation index (*F_st_*) analyses were conducted against comparator populations. *F_st_* values range from 0 (no allele frequency difference) to 1 (fixed difference). Unlike haplotype-based approaches, such as *iHS* or *XP-EHH*, which are most sensitive to recent selection events, *F_st_* can also capture older or historical signals of selection that may no longer present extended haplotypes due to recombination. This makes *F_st_* particularly valuable for detecting persistent, region-specific differentiation that reflects long-standing selective pressures within a population.

Our analyses focused primarily on loci potentially associated with historical drug pressure or recent hard sweeps unique to Ethiopia. Three SNPs exhibited extreme differentiation (*F_st_* ≥ 0.95) and were nearly fixed in Ethiopia while absent in other African countries with low, or no, endemic *P. vivax* (Table 1). The most differentiated variant, *pfcrt* N326S (*F_st_* = 0.997), is commonly linked to chloroquine resistance and may have epistatic interactions with pfk13 R622I; implicated in artemisinin resistance in northern Ethiopia and Southeast Asia^2,7,26^. Compared with all African countries in our dataset, regardless of *P. vivax* prevalence, *pfcrt* N326S remained the only SNP with a significant *F_st_* in Ethiopia (*F_st_* = 0.97).

**Table 1:**
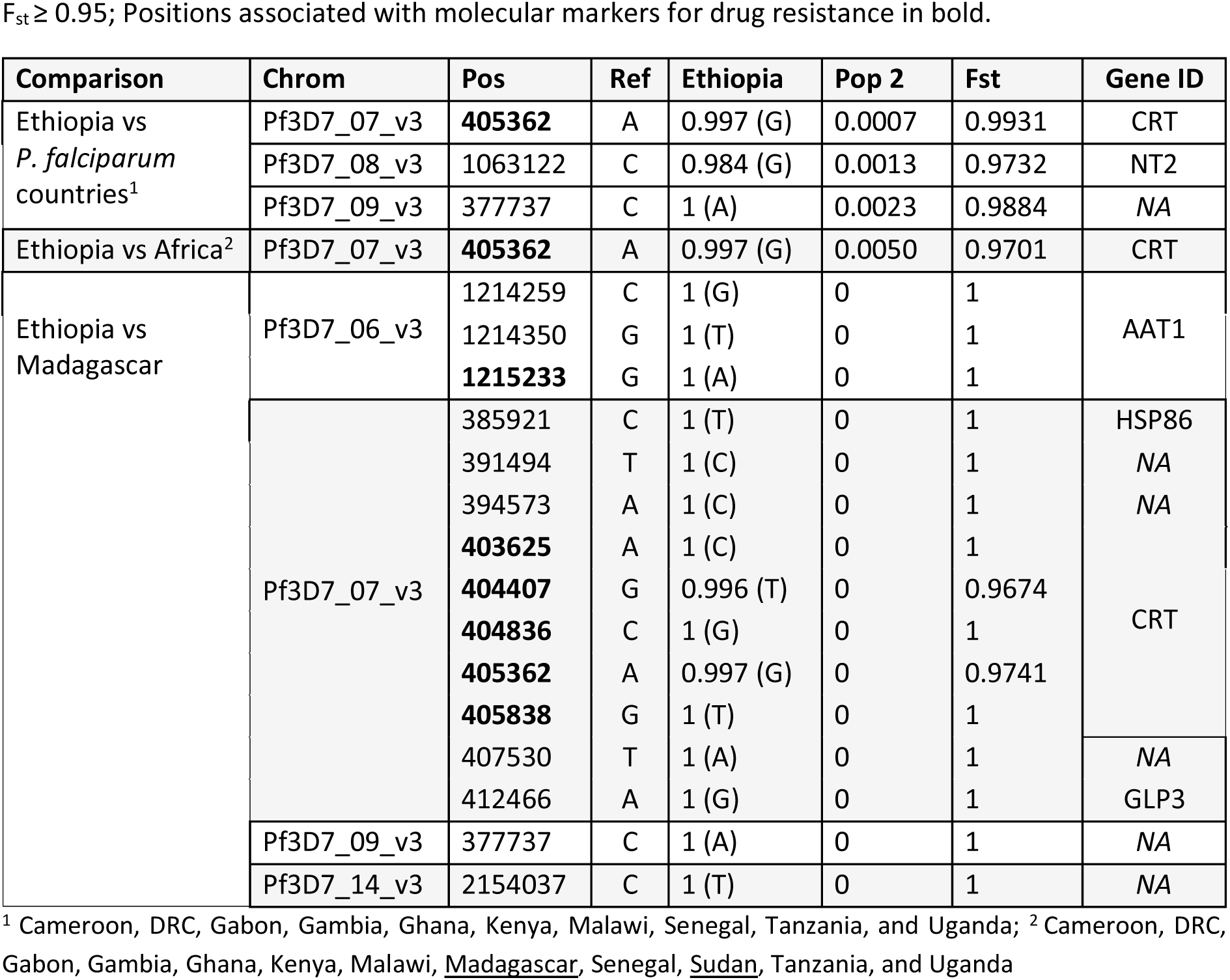
Significant Fst values for positions across the genome in Ethiopian and sub-Saharan African population. F_st_ ≥ 0.95; Positions associated with molecular markers for drug resistance in bold.

| Comparison | Chrom | Pos | Ref | Ethiopia | Pop 2 | Fst | Gene ID |
| --- | --- | --- | --- | --- | --- | --- | --- |
| Ethiopia vs *P. falciparum* countries<sup>1</sup> | Pf3D7_07_v3 | **405362** | A | 0.997 (G) | 0.0007 | 0.9931 | CRT |
|  | Pf3D7_08_v3 | 1063122 | C | 0.984 (G) | 0.0013 | 0.9732 | NT2 |
|  | Pf3D7_09_v3 | 377737 | C | 1 (A) | 0.0023 | 0.9884 | NA |
| Ethiopia vs Africa<sup>2</sup> | Pf3D7_07_v3 | **405362** | A | 0.997 (G) | 0.0050 | 0.9701 | CRT |
| Ethiopia vs Madagascar | Pf3D7_06_v3 | 1214259 | C | 1 (G) | 0 | 1 | AAT1 |
|  |  | 1214350 | G | 1 (T) | 0 | 1 |  |
|  |  | **1215233** | G | 1 (A) | 0 | 1 |  |
|  | Pf3D7_07_v3 | 385921 | C | 1 (T) | 0 | 1 | HSP86 |
|  |  | 391494 | T | 1 (C) | 0 | 1 | NA |
|  |  | 394573 | A | 1 (C) | 0 | 1 | NA |
|  |  | **403625** | A | 1 (C) | 0 | 1 | CRT |
|  |  | **404407** | G | 0.996 (T) | 0 | 0.9674 |  |
|  |  | **404836** | C | 1 (G) | 0 | 1 |  |
|  |  | **405362** | A | 0.997 (G) | 0 | 0.9741 |  |
|  |  | **405838** | G | 1 (T) | 0 | 1 |  |
|  |  | 407530 | T | 1 (A) | 0 | 1 | NA |
|  |  | 412466 | A | 1 (G) | 0 | 1 | GLP3 |
|  | Pf3D7_09_v3 | 377737 | C | 1 (A) | 0 | 1 | NA |
|  | Pf3D7_14_v3 | 2154037 | C | 1 (T) | 0 | 1 | NA |

<sup>1</sup> Cameroon, DRC, Gabon, Gambia, Ghana, Kenya, Malawi, Senegal, Tanzania, and Uganda; <sup>2</sup> Cameroon, DRC, Gabon, Gambia, Ghana, Kenya, Malawi, Madagascar, Senegal, Sudan, Tanzania, and Uganda

Comparisons between Ethiopia and Sudan, where *P. vivax* is also highly prevalent, revealed no significantly differentiated SNPs. The geographical proximity of these populations may limit population differentiation. In 2017, Sudan phased out the use of chloroquine for treatment of uncomplicated *P. vivax* in preference of AL+PQ, the *pfcrt* K76T variant, associated with chloroquine resistance, was present in the Sudanese population at the point of cessation (43%; 13/29) and future surveillance will be needed to monitor this trend^38^. This pattern likely reflects historical chloroquine exposure and shared selective histories rather than ongoing chloroquine pressure.

Madagascar provides a distinct contrast since although *P. vivax* is also highly prevalent, chloroquine was discontinued almost 20 years ago when the country transitioned to AL as the first-line treatment for all malaria species^39^. Here, 15 SNPs across four chromosomes displayed extreme differentiation (*F_st_*≥ 0.95) relative to Ethiopian isolates. Three of these were located within *pfaat1*, including the S258L variant (*F_st_* = 1), a known potentiator of chloroquine resistance. Several additional highly differentiated positions were identified within *pfcrt*, including K76T, A220S, Q271E, N326S, and R371I, highlighting the persistence of multi-site selective signatures on this locus

### Admixture analysis highlights divergent origins of Ethiopian P. falciparum populations

Admixture analysis using genome-wide SNPs from *P. falciparum* populations across sub-Saharan Africa (861,415 SNPs; N = 1,017) identified six major ancestral components (K = 6) that broadly aligned with geographic subregions (Supp. Fig. S10A). Genetic clusters corresponded closely to geography, with ancestral and transmission connectivity reflecting regional proximity (Fig. 4A). The six subregional groupings were defined as Central Africa ([CAF] Cameroon, Gabon; N = 155), East Africa ([EAF] Kenya, Tanzania, Uganda; N = 210), Horn of Africa ([HAF] Ethiopia, Sudan; N = 216), South-Central Africa ([SCAF] Democratic Republic of Congo; N = 118), Southeast Africa ([SEAF] Madagascar, Malawi; N = 121), and West Africa ([WAF] Gambia, Ghana, Senegal; N = 197)^8^.

**Figure 4:**
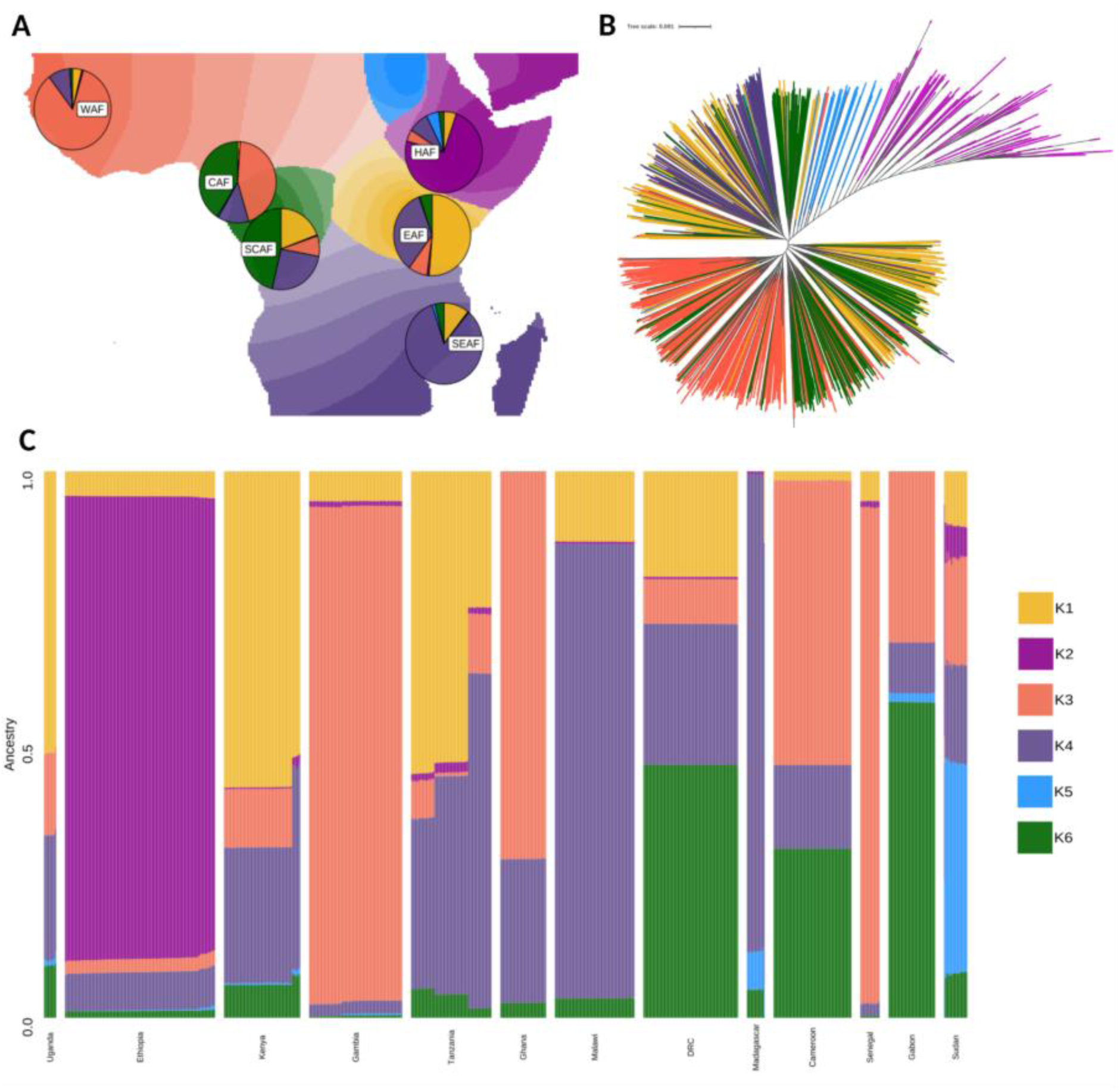
Ancestral admixture analysis of Ethiopian and sub-Saharan African *P. falciparum* **populations (N = 1,017; SNPs = 861,415).** (A) Geographic map of ancestry coefficients K1 to K6 with pie charts per region including all donor coefficients. (B) Maximum likelihood, unrooted, tree of isolates used to calculate admixture coefficients with isolates (nodes) coloured according to their predominant ancestry proportion (K) [N = 1,017; SNPs = 952,247]. (C) Bar plot of ancestry proportions per isolate (rows) by country (columns).

Despite their geographic proximity, Ethiopian and Sudanese *P. falciparum* parasites exhibited distinct ancestry signatures. Ethiopian isolates were overwhelmingly composed of the K2 ancestral component (84.6%), whereas Sudanese isolates were dominated by K5 (38.2%) and contained only a small fraction of K2 (7.2%) (Fig. 4A–B; Supp. Fig. S10B–C). An unrooted maximum-likelihood tree and IBD network analysis further supported the genetic distinctiveness of Ethiopian and Sudanese populations (Fig. 4B; Supplementary Fig. S10D). Both K2 and K5 ancestries were rare elsewhere in Africa (<7%), reinforcing the unique evolutionary trajectories of the Horn of Africa parasite lineages and suggesting that Ethiopian *P. falciparum* populations are genetically isolated reservoirs shaped by local drug and ecological pressures rather than continental gene flow.

## Conclusion

Our genomic analysis reveals a *Plasmodium falciparum* population in southern Ethiopia that is highly structured and genetically constrained, likely shaped by atypical local selection pressures. The continued fixation of *pfcrt, pfdhfr,* and *pfdhps* resistance haplotypes despite discontinued use of chloroquine and SP for treatment of *P. falciparum*, together with the emergence of a novel *pfmdr1* promoter deletion, highlights the resilience and adaptability of these parasites. Although partial artemisinin resistance associated with *pfk13* variants have yet to be confirmed in Southern Ethiopia, strong connectivity to northern Ethiopia, and signals of selection at *pfcrt* and *pfmdr1*, indicate a permissive genetic background that threaten the future spread of resistance and novel emergence in this region.

Our findings underscore how the unique epidemiological context of the Horn of Africa, marked by low transmission, limited recombination, and *P. vivax* co-endemicity, has produced an evolutionary trajectory distinct from that of most other African *P. falciparum* populations. Sustained genomic surveillance, tailored to local ecology and treatment practices, will be essential to anticipate resistance evolution and guide effective malaria elimination strategies in this region.

## METHODS

### Study site and population characteristics

Arba Minch is a city within the South Ethiopia Regional State with a tropical savanna climate conducive to year-round malaria transmission. *P. falciparum* and *P. vivax* are sympatric in this region, uncommon within most sub-Saharan countries. The transmission of *P. vivax* is largely attributed to the relative absence of the protective Duffy-negative phenotype in the population. Parasite isolates were collected as part of two clinical trials conducted in 2017 and 2021 (Clinical Trial IDs NCT01814683 and NCT03916003)^9,10^.

### Species identification and Sequencing

DNA extraction and speciation were carried out at Menzies School of Health Research, Australia. *Plasmodium* molecular speciation was performed using a multiplex PCR protocol, with modifications to the singleplex reaction for each species as recommended by Boonma *et al.* (2007)^40,41^.

Whole genome sequencing was performed at the Australian Genome Research Facility (AGRF), AU, and Sanger Institute, UK, on the Illumina NovaSeq platform using the Illumina DNA Prep M Library Prep kit and NovaSeq X 10B lane, 300 cycle (150 bp) sequencing kit (Illumina, USA). Samples with poor sequencing quality, missingness ≥70% with a minimum read depth of 5, were re-sequenced following *P. falciparum* sWGA^42^.

### Bioinformatic processing

The raw paired-end Illumina reads were trimmed using the VivaxGEN ngs-pipeline (GitHub: https://github.com/vivaxgen/ngs-pipeline) the *fastp* software suite was used prior to mapping. Mapping was done using *bwa-mem2*^43–47^. For genome-wide analyses, samples with less than 70% of reads mapped, and less than 5-fold coverage across 70% of the genome, were excluded. Variant calling was performed using *GATK4 - classic mode* (non-DRAGEN) with variant validation and quality control carried out using *GATK HaplotypeCaller, GenotypeGVCFs,* and *ApplyBQSR*; non bi-allelic SNPs were also removed from genome-wide analyses^48,49^. Of the 222 Ethiopian samples that were sequenced, 163 were suitable for use in genome-wide analyses, including 34 samples collected in 2017 and 129 samples collected in 2021. These were combined with 24 Ethiopian isolates from the Pf7 data release collected in 2013. Our Ethiopian isolates were combined with publicly available sub-Saharan African samples from the MalariaGEN Pf7 data release^50^. These samples were processed following the above methodology prior to being included in genome-wide analyses.

### Assessing drug resistance markers and structural variants

Structural variants (SVs) in *pfhrp2, pfhrp3*, and *pfmdr1* were identified using the Integrative Genomics Viewer (IGV) platform and custom Python-based workflows.

For looking at focal deletions and signals of larger structural variants, the Python-based workflow assessed paired-end BAM files by scanning across the regions of interest, as well as a their flanking regions, in 100 bp windows to quantify read-pair orientation, coverage loss, and small deletion events (20–80 bp)^32,51,52,31,50,51^. Small indel events on *pfmdr1* were identified by parsing CIGAR strings for deletion (D) operations of 20–80 bp among primary read alignments with mapping quality ≥30.

Indel events were aggregated by genomic position and deletion length. A deletion was considered present in a sample if ≥5 deletion-supporting reads originating from ≥5 independent read start positions were observed. For paired-end data, each template was counted once. Indel prevalence at the population-level was calculated using only samples meeting coverage requirements. Prevalence among Ethiopian isolates was compared with the African dataset using identical criteria.

For *pfhrp2* and *pfhrp3*, per-base read depth across the target region and flanking regions was calculated from aligned reads. Flanking region length varied depending on target gene due variable breakpoint positions, with some *pfhrp3* breakpoints occurring as far away as ∼30,000 base pairs from the gene position depending on the region. For automated coverage-based variant detection, samples with ≥10 median coverage, and a 20x coverage ≥70%, in both the right and left flanking regions were scanned for deletions. All samples were validated using manual IGV inspection in addition automated coverage and depth checking.

### Genetic measures of population diversity and dynamics

Identity-by-descent (IBD) was used to assess both connectivity and genome conservation between parasite populations, as well as a proxy for genetic selection within and between populations. IBD fractions were calculated by measuring the pairwise shared ancestry of genomic segments with these fractions representing parts of the genome that were inherited, without undergoing recombination events, from a recent common ancestor. Only SNPs with a minor allele frequency (MAF) of at least 0.01 were retained while the minimum genomic region length, or window size, was set to 50kb and the quantile cutoff for significance was 0.90. The software *hmmIBD* was implemented to estimate IBD while accounting for recombination using a hidden Markov model-based approach^53^.

To identify signals of directional selection and evidence of strong selective sweeps, the *rehh* R-based software was used to calculate haplotype heterozygosity within populations, the integrated haplotype score (*iHS*) as well as between populations, extended haplotype heterozygosity (XP-EHH)^54^. SNPs in regions of the genome exhibiting an *iHS* value of ≥2.5 or ≤-2.5 were classified as significant^55,56^.

Positive *iHS* scores are associated with haplotypes carrying ancestral alleles that are longer than the new allele, indicating the ancestral allele is under selection. A negative *iHS* score is associated with a new and rapidly increasing allele located on a haplotype longer than the ancestral allele and indicates the newly derived allele is under selection^54,56^. The same significance cutoff was applied to XP-EHH values, with values ≥ 2.5 and ≤ -2.5 considered significant. SNPs and samples with a missingness ≥ 30% were not included in calculations. Weir and Cockerham’s *F_st_* was calculated using an in-house R script. *F_s_* positions with missingness of ≥ 80% not included in calculations.

To estimate multiplicity of infection, *F_ws_* metrics were generated using the R-based package *moimix*^57^. Approximately 50.6% of *P. falciparum* isolates were polyclonal (*F_ws_* ≤ 0.95; N = 515), while 49.4% were monoclonal (*F_ws_*≥ 0.95; N = 502). For tree generation and admixture analyses, heterozygous SNP loci were converted to homozygous calls by assigning the major genotype (861,415 SNPs). Major alleles were determined at each polymorphic locus using allele depth (AD) where a major allele must have an AD ≥52%. A position with an AD too close to 50% was assigned as missing.

A maximum likelihood phylogenetic tree was generated using *IQTree* ModelFinder Plus (MFP) algorithm which automatically tested and selected the best-fit nucleotide substitution model^58^. Of the 610,354 SNPs used to inform the algorithm, 220,603 were parsimony-informative sites while 386,751 were singleton sites. There were no constant sites in the dataset. The model of best fit was a fully flexible substitution model using observed base frequencies that corrected for missing sites and allowed for a flexible variation rate across sites to handle high rates of heterogeneity within the dataset. The tree was annotated using the *iTOL* web interface^59^.

To determine population structure and ancestry of isolates, model-based ancestry coefficients were generated using *tess3r*^60,61^. *tess3r* combines matrix factorization methods and spatial statistics to provide estimates of ancestry matrices based on least-squares approximation. Root mean-squared errors were computed on a subset of loci used for cross-validation of the K value of best fit (ALPS; method =”projected.Is”). An optimum K value was estimated to be 6 using cross entropy values generated from dimensions 1 to 10 and based on previously established genetic clusters in Africa6. The tess3r algorithm for K value equals 6 was run 50 times using default parameters, including a spatial regularization parameter (σ = 1) to balance low and penalty functions, and the best 𝑄-matrix of ancestry coefficients for each isolate was retained. The major ancestral groupings were defined as K1 (Kenya, Tanzania, Uganda), K2 (Ethiopia), K3 (Cameroon, Gambia, Ghana, Senegal), K4 (Southeast Africa; Malawi, Madagascar), K5 (Sudan), K6 (Democratic Republic of Congo, Gabon).

To assess connectivity between *P. falciparum* isolates from Ethiopia that carry the R622I pfk13 variant and our isolates from Arba Minch, publicly available Molecular Inversion Probe (MIP) data from sites across Ethiopia with mutant lineages was combined with the WGS data from Arba Minch^5^. The WGS data was subset to the genome-wide MIP positions. Samples with less than 80% of positions with a minimum read depth of 5 were excluded from these analyses. A final dataset of 525 samples (114 WGS isolates from Arba Minch) was used to assess connectivity between populations. Connectivity was determined by calculating IBD across 1,061 of the MIP positions using hmmIBD^53^. MIP positions that did not pass the filtering criteria were not included (i.e. ≥80% of positions with a minimum read depth of 5).

### Ethical approval

All samples in this study were derived from blood samples obtained from patients positive for malaria and collected with informed consent from the patient, or patient parent / legal guardian where individuals were less than 18 years of age. At each location, sample collection was approved by the appropriate local institutional ethics committees. The following committees gave ethical approval for the partner studies; Human Research Ethics Committee of NT Department of Health and Families and Menzies School of Health Research, Darwin, Australia (HREC-2013-1991 and HREC-2019-3288); the National Research Ethics Review Committee of Ethiopia (Ref No. EPHI6.13/4362009); College of Medicine and Health Sciences Institutional Research Ethics Review Board, Arbaminch, Ethiopia (CMHS/121733883/111); Columbia University Medical Center Institutional Review Board, New York, United States (IRB-AAAQ3503);

## Code availability

The *VivaxGEN* ngs-pipeline for retrieving SNP-based variant calls is available on GitHub: https://github.com/vivaxgen/ngs-pipeline. A bespoke GitHub repo for all downstream population genetic analyses, *malaria-popgen-toolkit,* is available at https://github.com/aosborne13/malaria-popgen-toolkit.

## Data availability

The MalariaGEN Pf7 genomic data are available through the European Nucleotide Archive^50^. The genomic reads from the newly generated, high-quality, Ethiopian *P. falciparum* infections will be made available in the European Nucleotide Archive at publication.

## Acknowledgements

This work was supported by the National Health and Medical Research Council of Australia (Investigator Grant 2025377 to S.A.), the Medical Research Council and UK Department for International Development (award number M006212 to D.K.) and the Wellcome Trust (award numbers 206194 and 204911 to D.K.). We thank the patients who contributed their samples to the study, and the health workers and field teams who assisted with the sample collections. Whole genome sequencing was undertaken at the Australian Genome Research Facility, Australia, and the Wellcome Sanger Institute, UK.

## Author contributions

A.A.O. and S.A. conceived and designed the study. D.T., A. Rai and A. Rumaseb contributed to the laboratory coordination and quality assurance. A.A.O, H.T., C.L., R.D.P. and D.P.K. contributed whole genome sequencing data production and informatic support. A.A.O, H.T., C.L. and R.D.P contributed to bioinformatic pipeline development. A.O. conducted data analysis. A.A., T.S.D, A.H., A.W., D.T., R.N.P. and K.T. contributed essential field-based malaria collections and metadata, and guidance on the study design and interpretation.

A.A.O. wrote the first draft of the manuscript, and all authors read and reviewed the manuscript draft. This manuscript has not been accepted or published elsewhere.

---

**Supplementary Figure S1.**
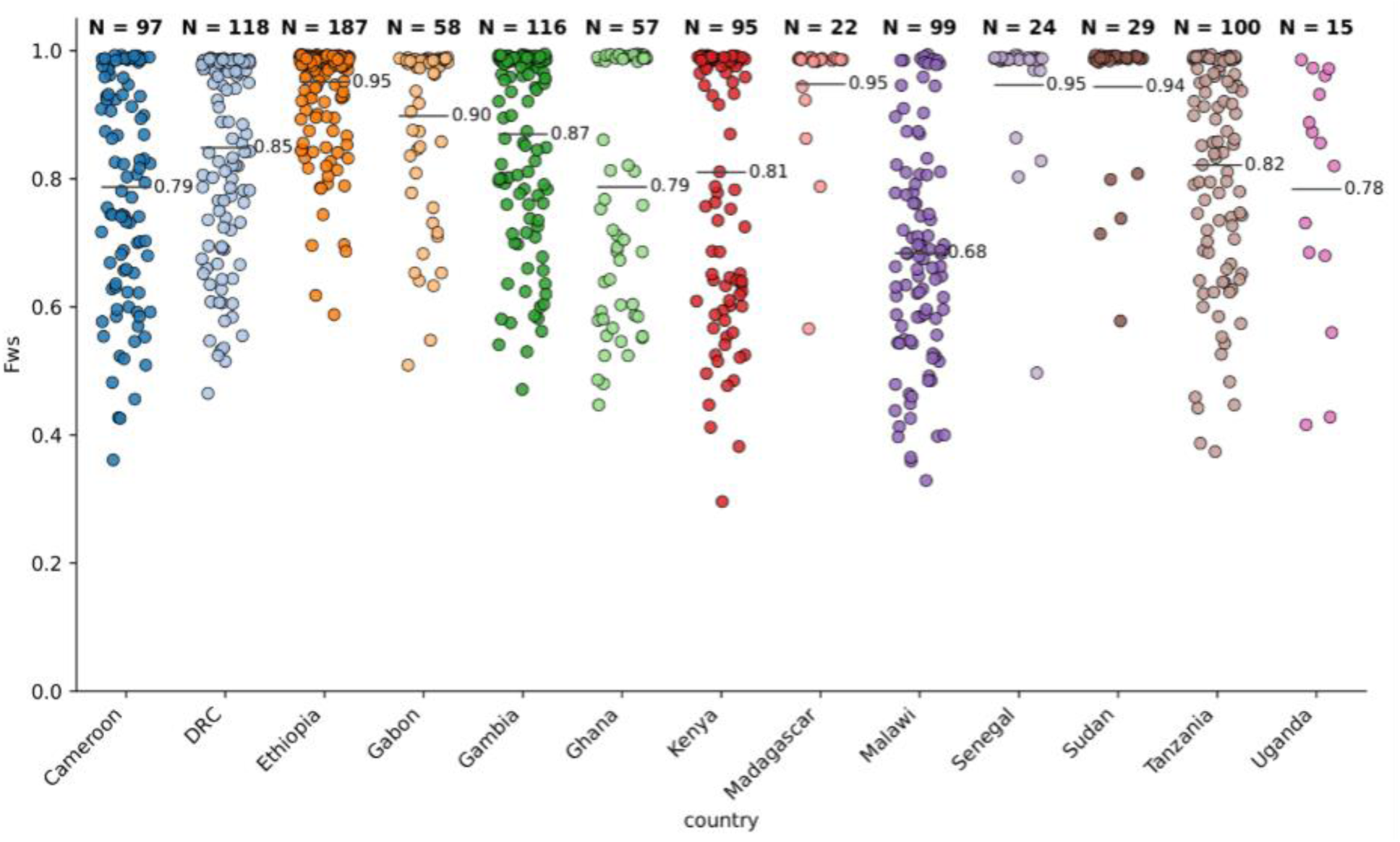
Multiplicity of infection calculated using the inbreeding coefficient (F_ws_ metric; R-package MOImix). Samples by country included in the African-wide population-level analyses (N = 1,017).

**Supplementary Figure S2:**
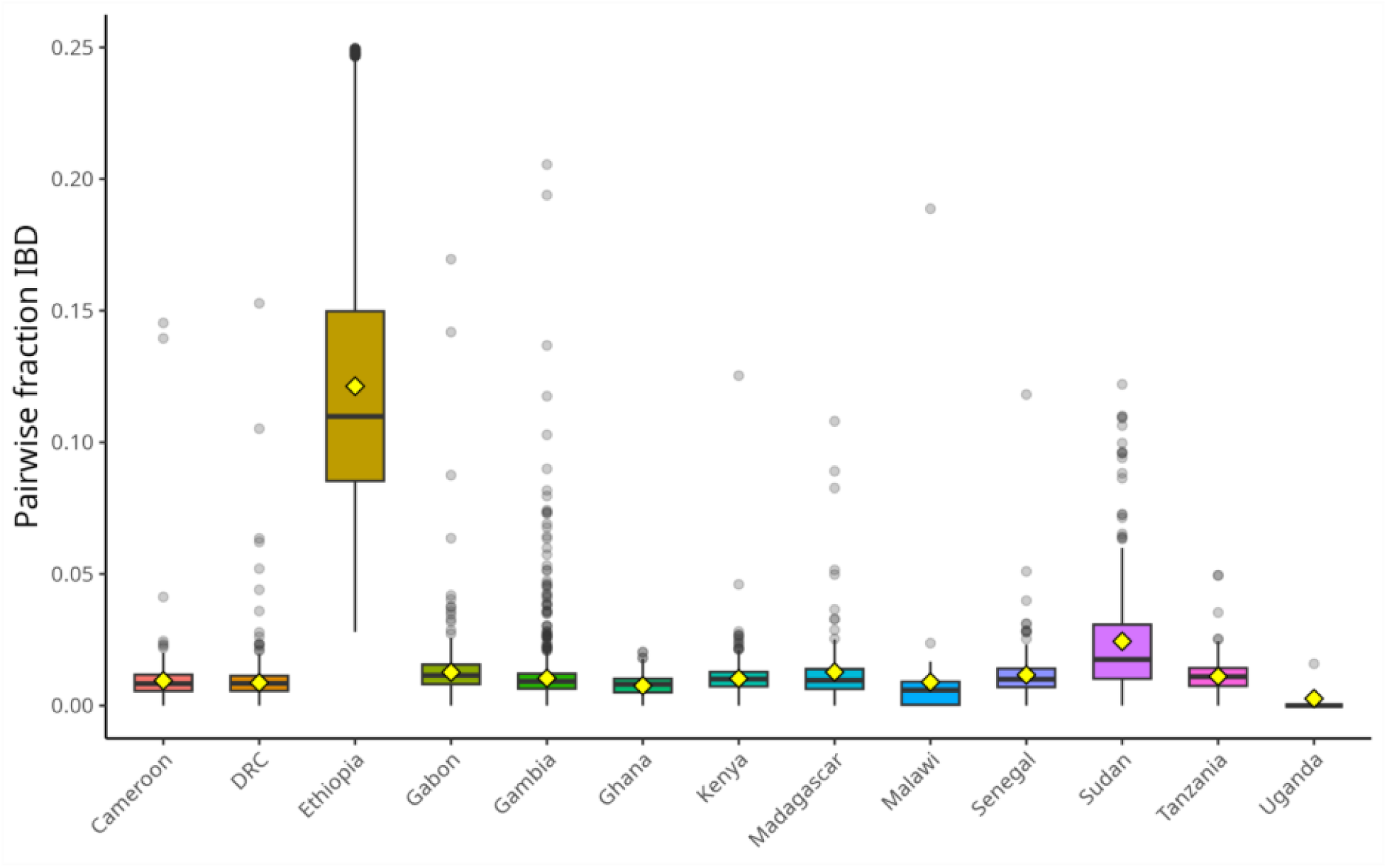
Pairwise fractions of Identity-by-Descent within *P. falciparum* populations according to country (F_ws_ ≥0.95; N = 502). Distribution is the mean IBD of all pairwise comparisons of samples by country.

**Supplementary Figure S3:**
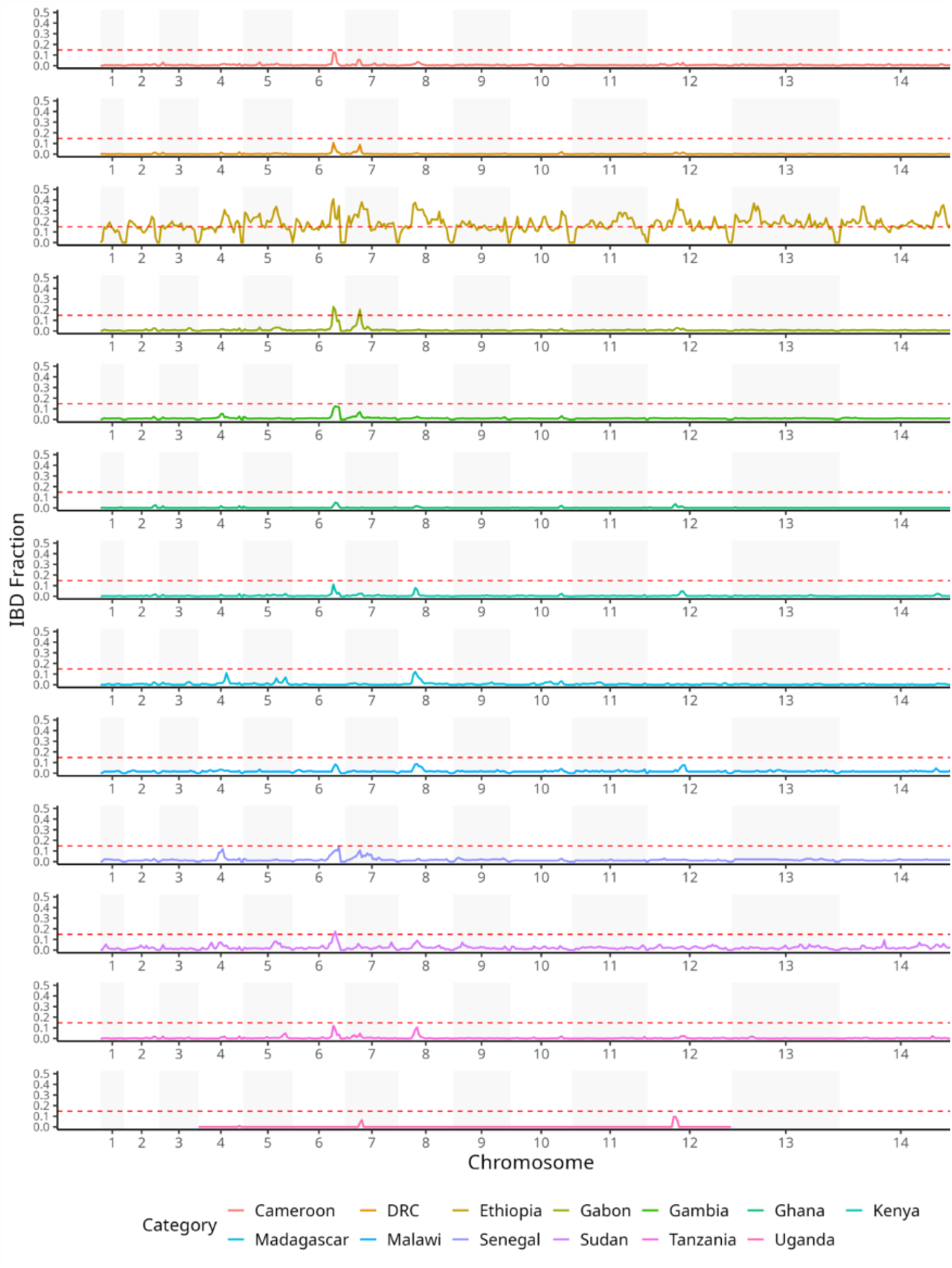
Identity-by-Descent fractions across the genome by country (N = 502; *F_ws_* ≥ 0.95).

**Supplementary Figure S4:**
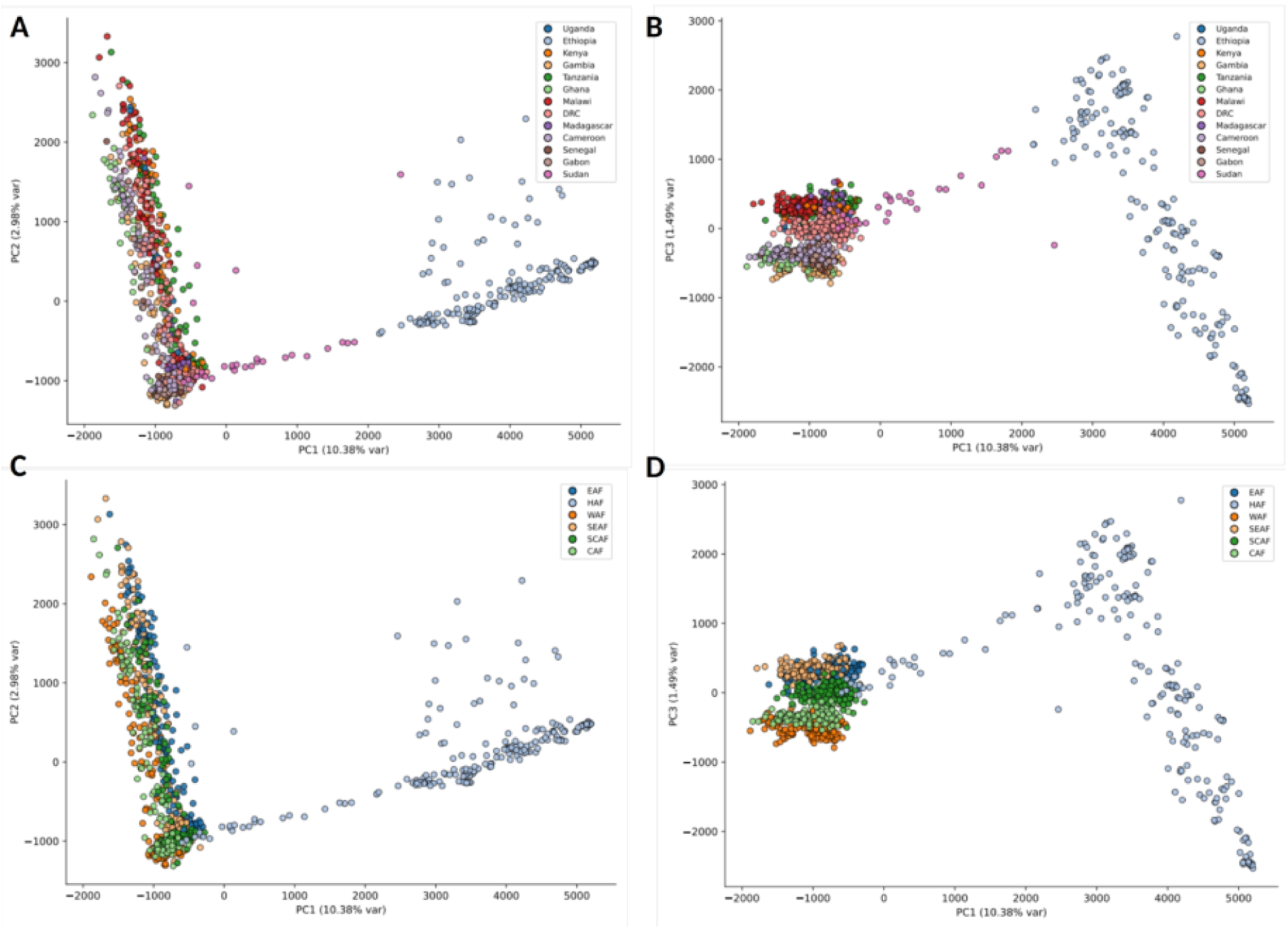
Principal component analysis (PCA) of all African isolates included in genome-wide analyses (A and B) Principal components 1, 2, and 3 showing differentiation by samples coloured by geographical region* within sub-Saharan Africa (N = 1,017; SNPs = 861,415). (C and D) Principal components 1, 2, and 3 showing differentiation by samples coloured by country. *CAF = Cameroon, Gabon; EAF = Kenya, Tanzania, Uganda; HAF = Ethiopia, Sudan; SCAF = Democratic Republic of Congo; SEAF = Madagascar, Malawi; WAF = Gambia, Ghana, Senegal

**Supplementary Figure S5:**
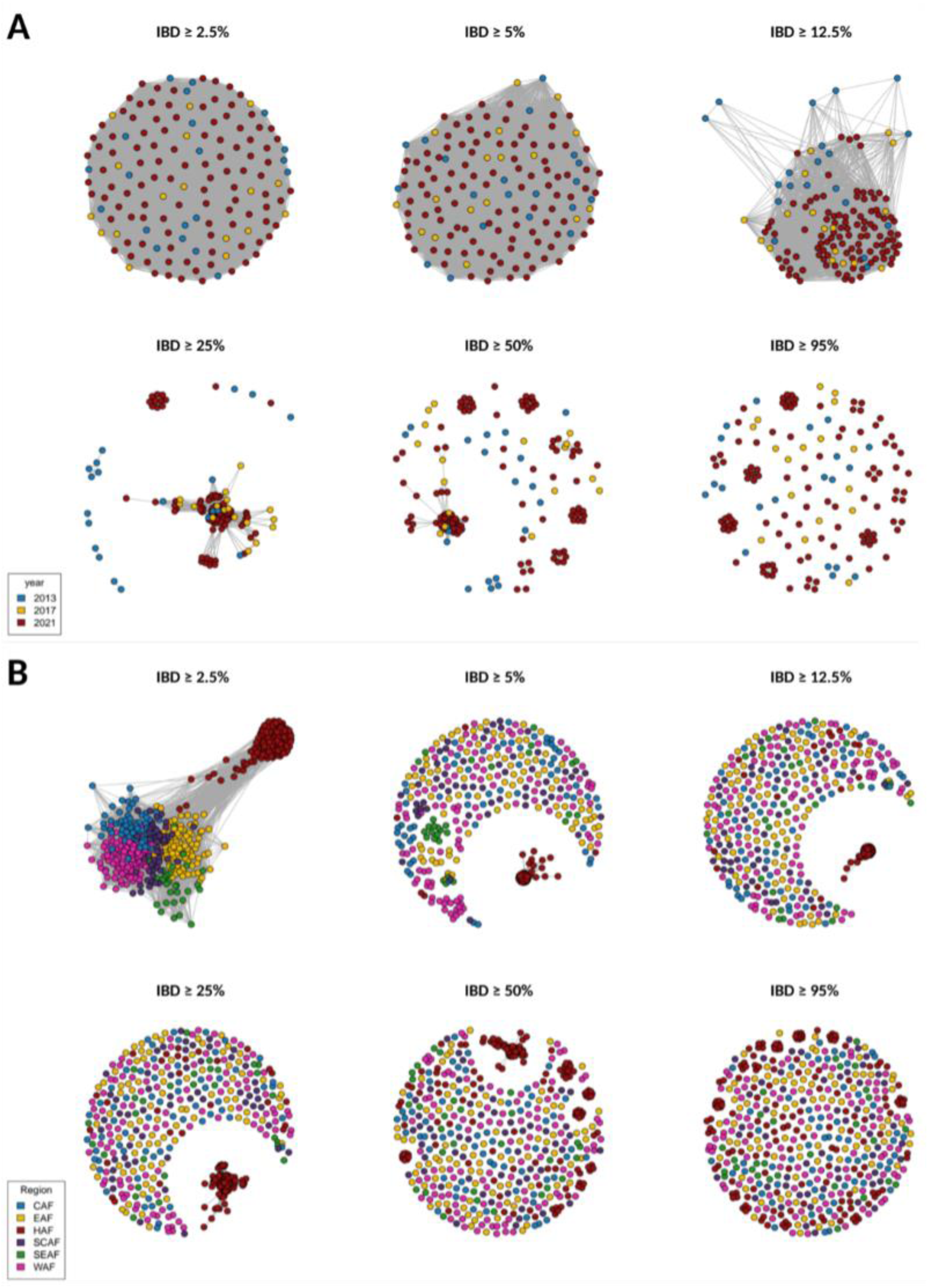
Connectivity between *P. falciparum* populations according to Identity-by-Descent thresholds. (A) IBD within Ethiopia over time (2013, N = 20; 2017, N = 17; 2021 = 105 [*F_ws_* ≥ 0.95]). (B) IBD between populations across the African continent defined by geographical region* (N = 502; *F_ws_*≥ 0.95). *CAF = Cameroon, Gabon; EAF = Kenya, Tanzania, Uganda; HAF = Ethiopia, Sudan; SCAF = Democratic Republic of Congo; SEAF = Madagascar, Malawi; WAF = Gambia, Ghana, Senegal

**Supplementary Figure S6:**
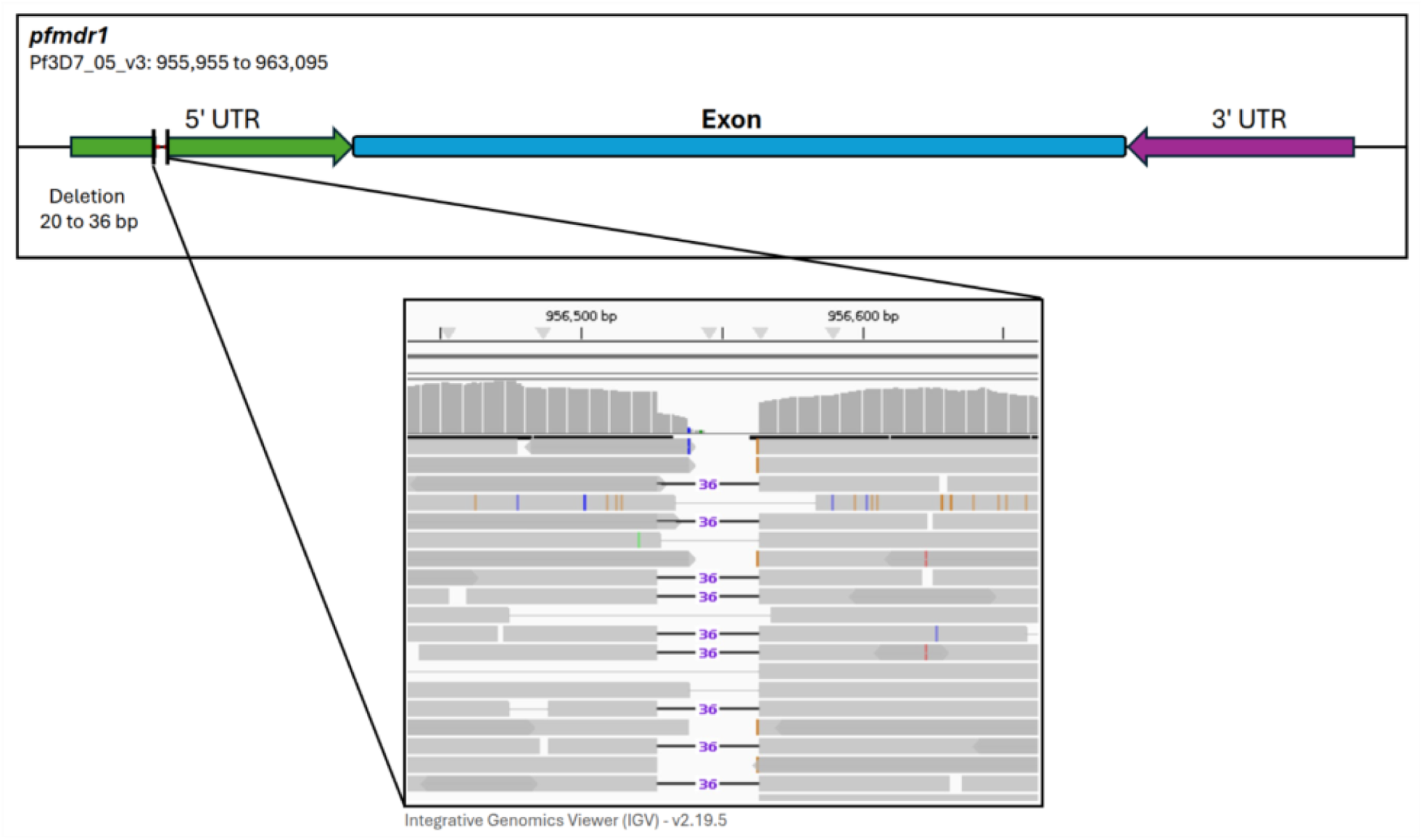
Schematic of *pfmdr1* 5’ untranslated region (UTR) indel observed in *P. falciparum* populations across sub-Saharan Africa. Read pair schematic rendered using IGV v2.19.5.

**Supplementary Figure S7:**
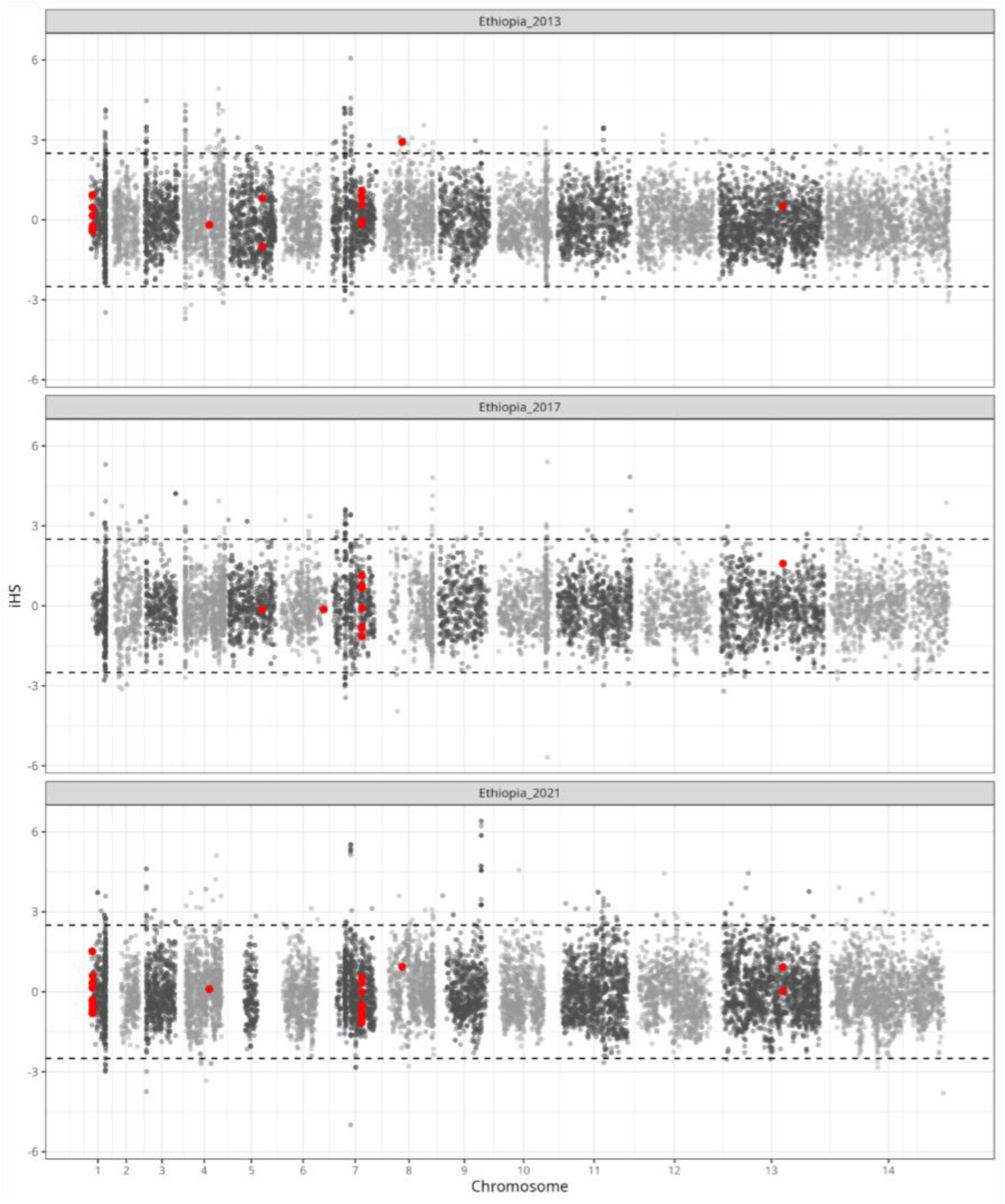
Signatures of selection in Ethiopian isolates by year of collection*. Genome-wide haplotype structure of isolates highlighting regions of high homozygosity compared with neutral expectation (*iHS*). Positions exhibiting an *iHS* value of ≥2.5 or ≤-2.5 classified as significant. Positions within drug resistance-associated genes highlighted in red (Supp. Table S2). *2013, N = 20; 2017, N = 17; 2021 = 105 (*F_ws_* ≥ 0.95)

**Supplementary Figure S8:**
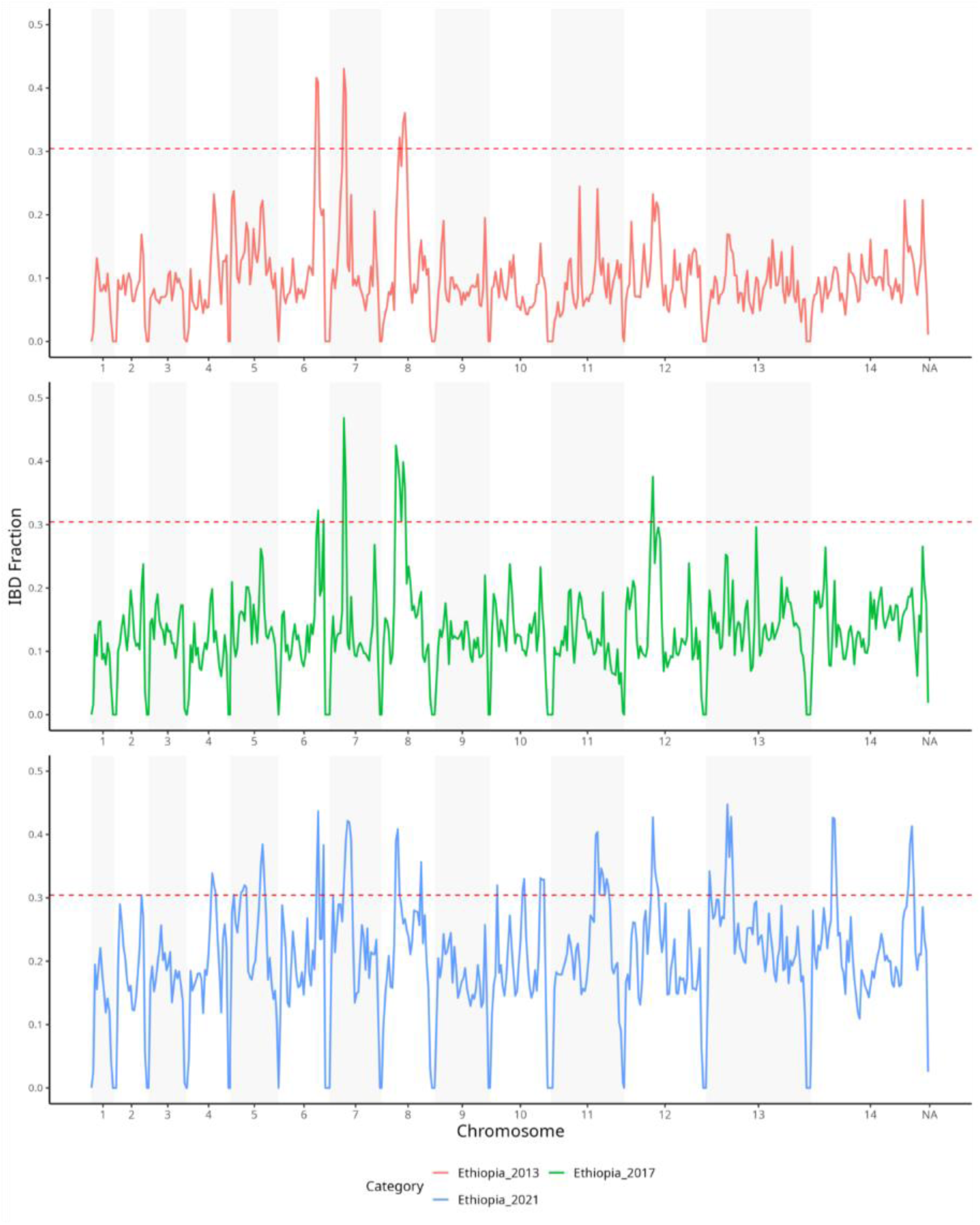
Genome-wide Identity-by-Descent fractions within Ethiopian populations striated by year of collection (2013, N = 20; 2017, N = 17; 2021 = 105 [*F_ws_*≥ 0.95]).

**Supplementary Figure S9:**
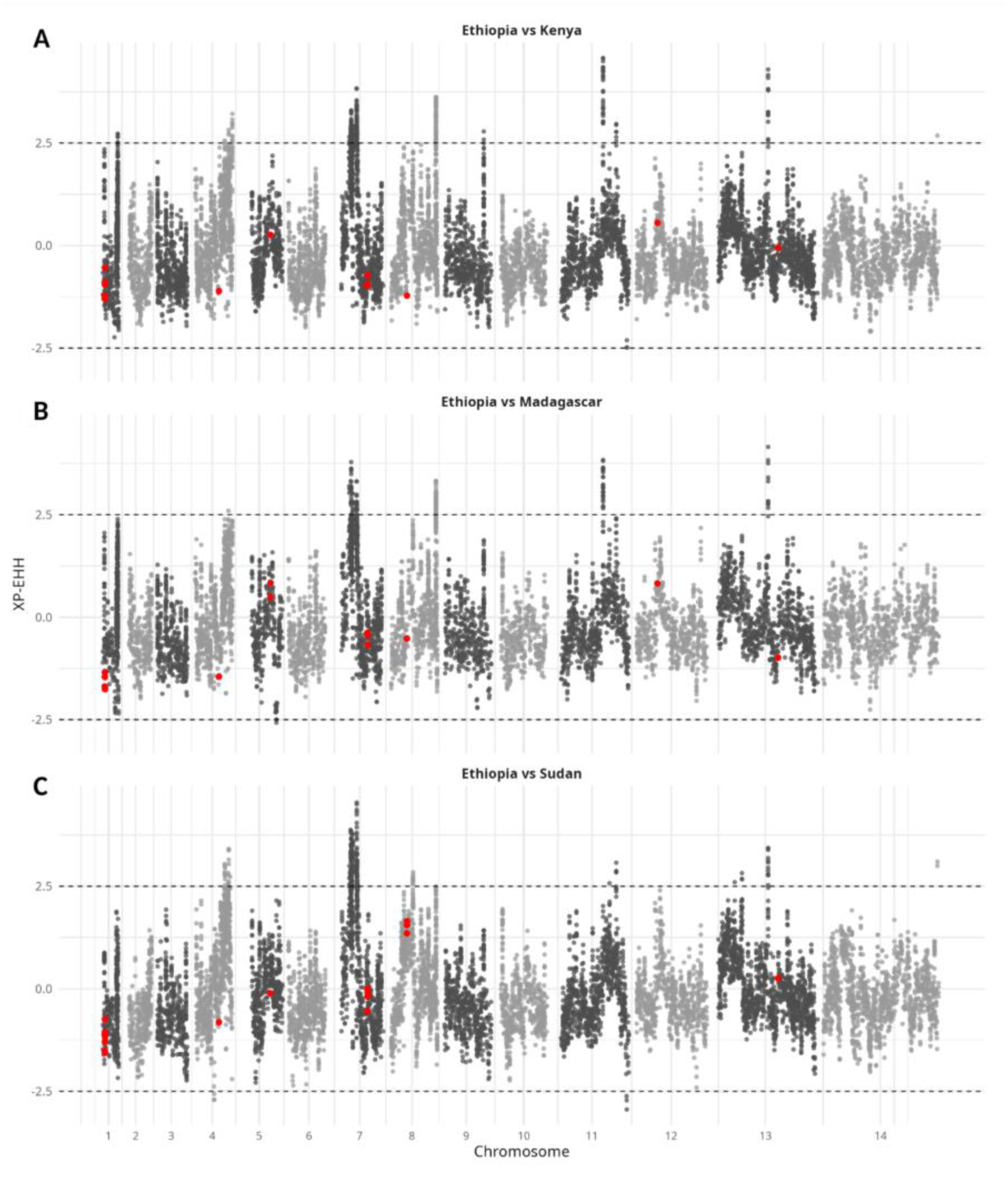
Cross-population analysis of variants exhibiting significant *XP-EHH* values between Ethiopian populations (N = 142; *F_ws_* ≥ 0.95) with: (A) Kenyan populations (N = 47; *F_ws_* ≥ 0.95); (B) Madagascan populations (N = 17; *F_ws_* ≥ 0.95); (C) Sudanese populations (N = 24; *F_ws_* ≥ 0.95). Positions within drug resistance-associated genes highlighted in red (Supp. Table S2)

**Supplementary Figure S10:**
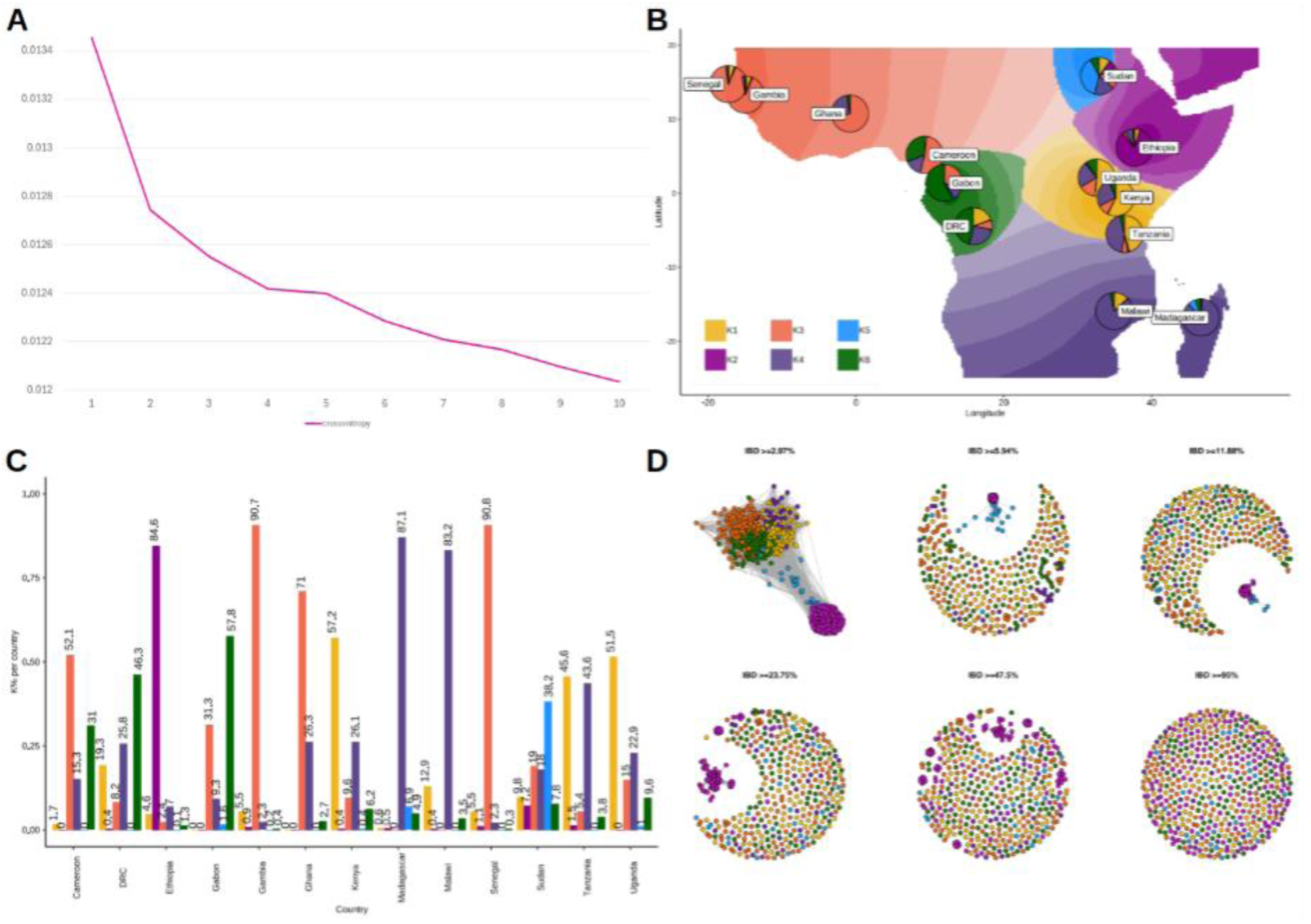
Ancestral admixture analysis of African subpopulations using the *Tess3r* software (N = 1,017; SNPs = 861,415). (A) Cross-validation estimate using entropy values for dimensions 1 to 10; (B) Geographic map of ancestry coefficients K1 to K6 with pie charts per country including donor coefficients; (C) Cumulative genome-wide ancestral admixture analysis proportions by country; (D) IBD network plot of African *P. falciparum* subpopulations coloured by predominant ancestry coefficient (K). The K1 ancestral population largely correlated with the EAF populations, including isolates from Kenya (K1 proportion; 57.2%), Tanzania (45.6%), and Uganda (51.5%). The second most predominant ancestral proportion in EAF was K4 with isolates from Tanzania exhibiting 43.6% of the K4 proportion, while Kenyan and Ugandan isolates contained 26.1% and 22.9%, respectively. The K4 ancestral population was strongly linked to SEAF with populations from Madagascar and Malawi having 87.1% and 83.2% of their ancestral proportions contributed by K4. The K3 ancestral population contributed large proportions to the WAF populations, including Gambia (K3 proportion; 90.7%), Ghana (71%), and Senegal (90.8%). Cameroon was also identified to share a high ancestral proportion with WAF populations (K3; 52.1%). Although classified as different geographic subregions, limited differentiation was observed between CAF and SCAF populations with Cameroon grouping with WAF populations and Gabon having a higher proportion of K6 ancestry (K6 proportion; 57.8%). The K6 ancestral population was split between Gabon and DRC (K6; 46.3%) with some contribution to Cameroonian populations (31%).

**Supplementary Table S1:** Summary data for 1,017 *P. falciparum* isolates included within African genome wide population analyses.

| Region | Country | Year | Samples |
| --- | --- | --- | --- |
| East Africa (EAF) | Kenya | 2014, 2015 | 95 |
|  | Tanzania | 2013 | 100 |
|  | Uganda | 2010, 2022 | 15 |
| Central Africa (CAF) | Cameroon | 2013, 2016, 2017 | 97 |
|  | Gabon | 2014 | 58 |
| Horn of Africa (HAF) | Ethiopia | 2013, 2017, 2021 | 187 |
|  | Sudan | 2015, 2017 | 29 |
| South Central Africa (SCAF) | Democratic Republic of Congo | 2015, 2016 | 118 |
| Southeast Africa (SEAF) | Madagascar | 2012, 2013, 2014 | 22 |
|  | Malawi | 2011 | 99 |
| West Africa (WAF) | Gambia | 2014 | 116 |
|  | Ghana | 2015 | 57 |
|  | Senegal | 2014 | 24 |

**Supplementary Table S2:**
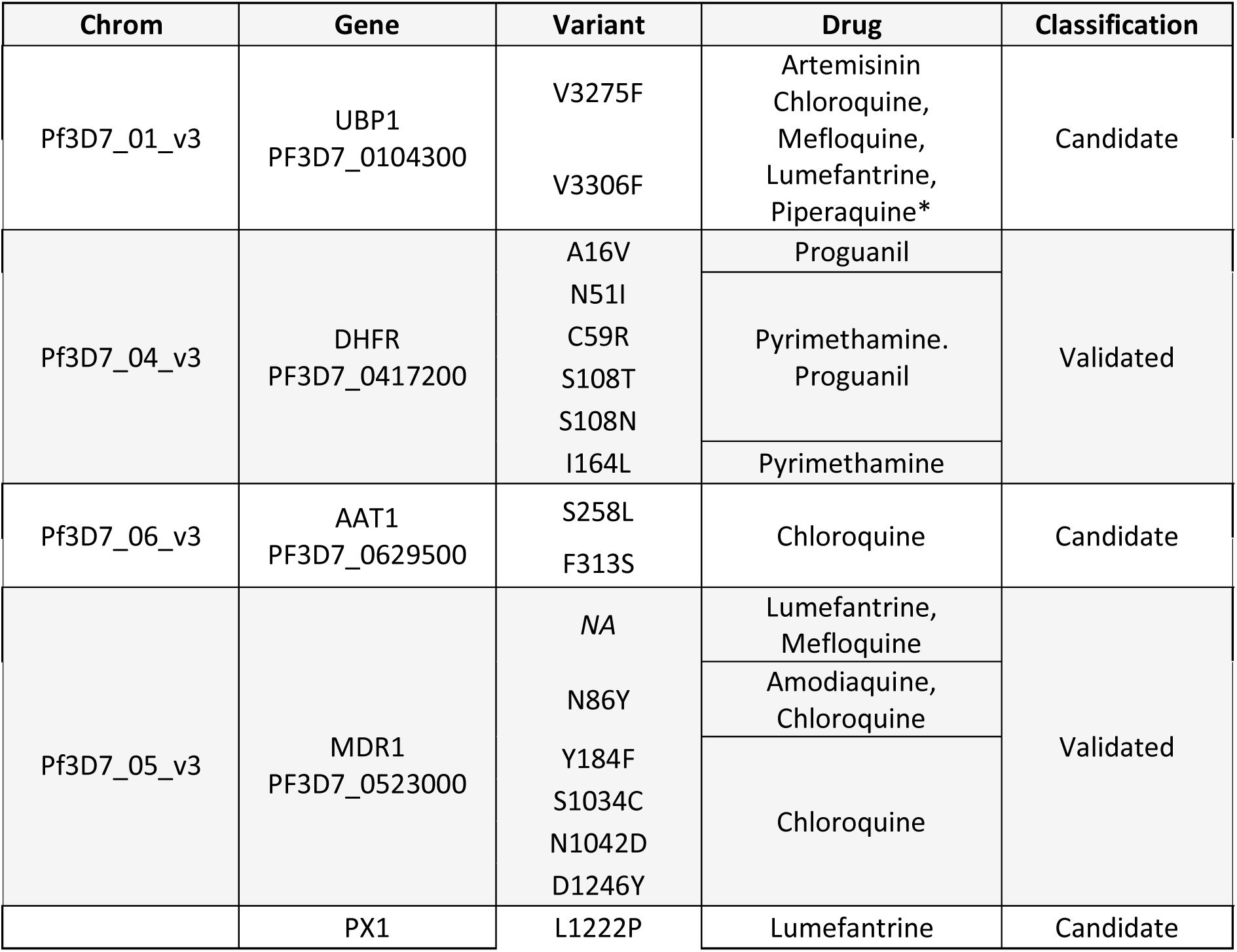
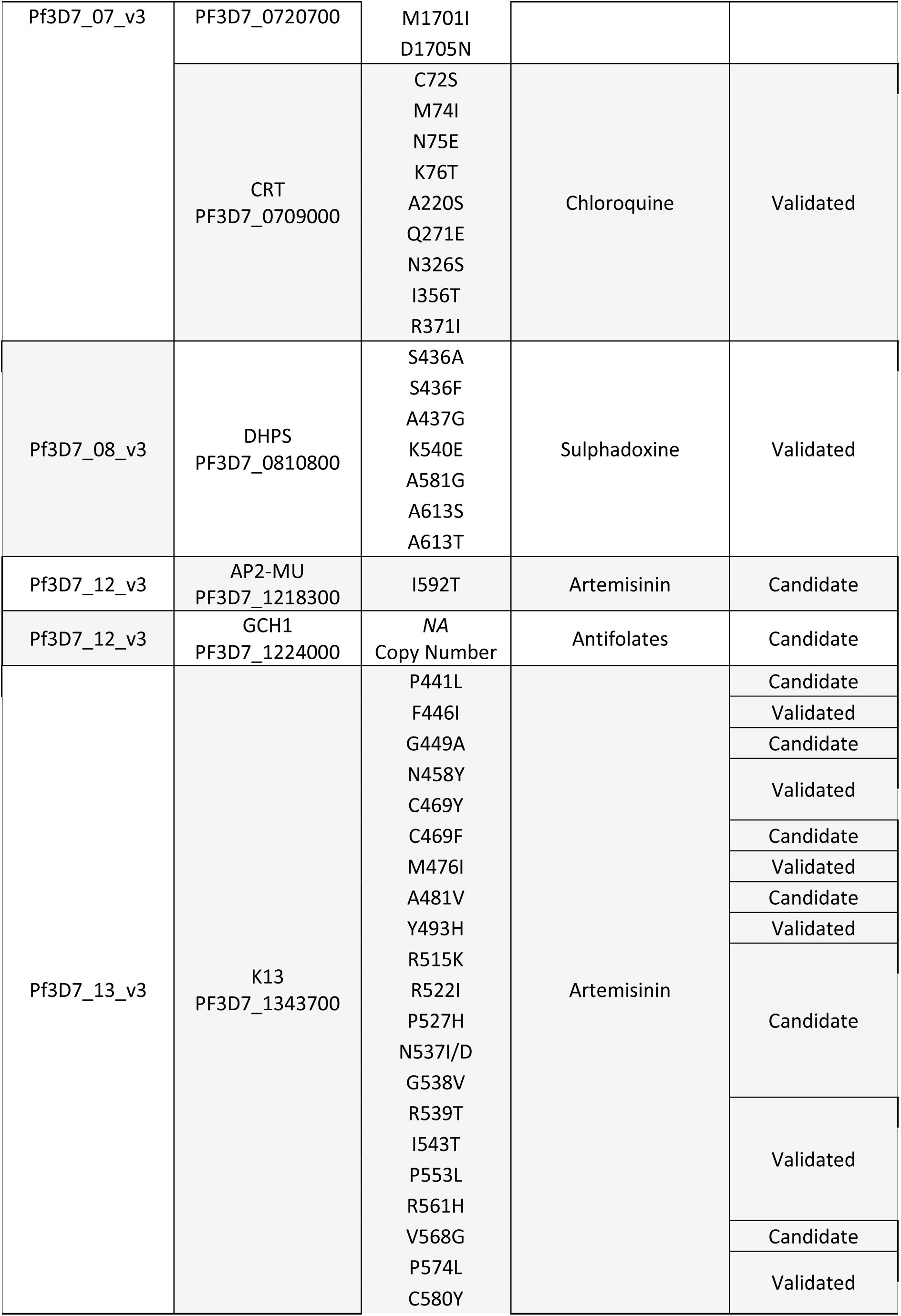
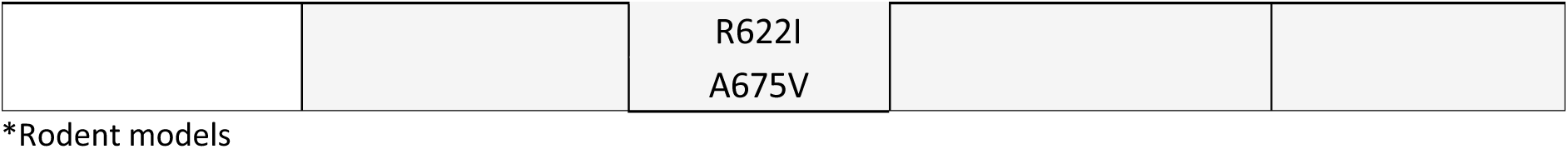
Candidate and validated molecular markers associated with drug resistance. Validated markers were characterized as requiring both laboratory and clinical support of the observed genetic alteration.

**Supplementary Table S3:**
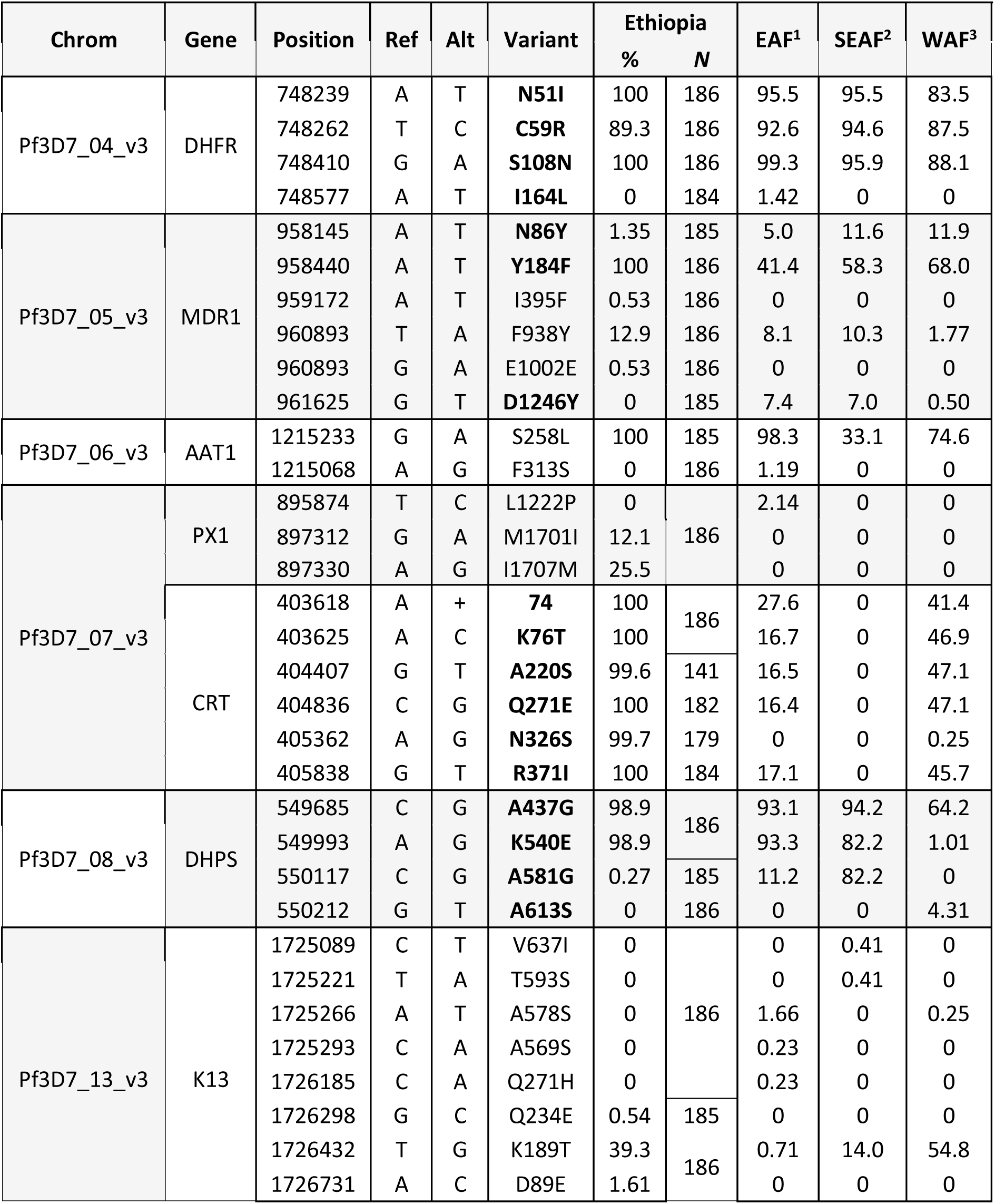
Proportions of non-synonymous polymorphisms (SNPs) within both candidate and validated drug resistance genes in Ethiopia and subpopulations of *P. falciparum* in sub-Saharan Africa. Validated molecular markers in Bold. Country breakdown in Supplementary Spreadsheet 1.

**Supplementary Table S4:** Proportion of *pfhrp2* and *pfhrp3* deletions in Ethiopian *P. falciparum* populations.

| Year | *pfhrp2* |  | *pfhrp3* |  |
| --- | --- | --- | --- | --- |
|  | % | N | % | N |
| 2021 | 7.41 | 8/108 | 56.4 | 57/101 |
| 2017 | 0 | 0/38 | 59.6 | 28/47 |
| 2013 | 0 | 0/20 | 42.9 | 9/21 |
| Total | 5.12 | 8/156 | 55.6 | 94/169 |

**Supplementary Table S5:** Proportion of *pfmdr1* 5’ UTR indel in Ethiopian and African *P. falciparum* populations. Median regional DP ≥25; ≥90% regional coverage at ≥20× DP.

| Country | | Population Frequency | | Proportion ($F_{ws} \geq 0.95$) | |
| --- | --- | --- | --- | --- | --- |
|  |  | % | N | % | N |
| Cameroon |  | 31.6 | 18/57 | 26.7 | 4/15 |
| DRC* |  | 66.6 | 2/3 | 50.0 | 1/2 |
| Ethiopia |  | 40.0 | 26/65 | 40.0 | 22/55 |
| Ethiopia | 2021 | 43.1 | 25/58 | 42.9 | 21/49 |
|  | 2017* | NA |  | NA |  |
|  | 2013* | 14.3 | 1/7 | 16.7 | 1/6 |
| Gabon |  | 34.3 | 12/35 | 20.0 | 4/20 |
| Gambia |  | 28.6 | 10/35 | 46.7 | 7/15 |
| Ghana* |  | NA |  | NA |  |
| Kenya |  | 40.0 | 14/35 | 39.1 | 9/23 |
| Madagascar |  | 38.5 | 5/13 | 36.4 | 4/11 |
| Malawi |  | 24.4 | 19/78 | 0 | 0/6 |
| Senegal* |  | 50.0 | 4/8 | 57.1 | 4/7 |
| Sudan* |  | NA |  | NA |  |
| Tanzania |  | 26.3 | 5/19 | 20.0 | 1/5 |
| Uganda* |  | 0 | 0/5 | 0 | 0/1 |

\*Poor read depth within *pfmdr1* 5' UTR

